# Time-resolved immune dynamics in rheumatoid arthritis under methotrexate therapy

**DOI:** 10.1101/2025.01.14.629357

**Authors:** Teresa Preglej, Anela Tosevska, Marie Brinkmann, Philipp Schatzlmaier, Elisabeth Simader, Daniela Sieghart, Philipp Hofer, Thomas Krausgruber, Lina Dobnikar, Christoph Bock, Thomas Karonitsch, Renate Kain, Hannes Stockinger, Wilfried Ellmeier, Daniel Aletaha, Lisa Göschl, Michael Bonelli

## Abstract

Rheumatoid arthritis (RA) is characterized by immune dysregulation, including alterations in peripheral blood mononuclear cell (PBMC) populations and aberrant cytokine signaling. Methotrexate (MTX) is the preferred first-line treatment for RA, yet its precise mechanisms of action remain incompletely understood. This study employed a multi-omics strategy—combining single-cell RNA sequencing (scRNA-seq) and immunophenotyping—to identify key effector peripheral immune cells and their cellular responses in RA patients over 12 weeks of MTX treatment.

In our study, MTX was associated with significant immune modulation, including the restoration of naïve T and B cells and reductions in T cell memory subsets with these effects detectable as early as three weeks post-treatment. Plasmablast levels also emerged as a potential biomarker for early therapeutic response, reflecting MTX’s impact on immune homeostasis. Transcriptional analysis revealed modulation of key pathways, including TNF-α signaling, B cell receptor signaling, and T cell receptor-mediated apoptosis. Network analysis identified critical regulatory hubs, such as EGR1, JAK2, and SOCS1, in monocytes and CD4 memory T cells, highlighting these cell types as key mediators of MTX’s effects.

In conclusion, these findings advance our understanding of MTX’s effects on immune cell dynamics at different stages of treatment, showing for the first time the early cellular changes leading to immune modulation in RA. Altogether, our results provide the foundation for further mechanistic investigations into MTX.

## Introduction

Rheumatoid arthritis (RA) represents a systemic autoimmune disease primarily affecting the joints and leading to cartilage degradation, bone erosion, and, if left untreated, irreversible destruction of the joints (1–4). While the precise etiology of RA remains elusive, the disease is marked by an aberrant immune cell activity that disrupts the delicate immunological balance (5–7). A complex interplay of diverse immune cell types contributes to the initiation and progression of RA. Among the key players are B cells, T cells, and myeloid cells, which infiltrate the synovium and circulate in peripheral blood, perpetuating inflammatory and destructive processes (5–7).

Inadequately treated RA results in severe joint damage, disability, and other systemic manifestations (1–4). The advent of disease-modifying anti-rheumatic drugs (DMARDs) has revolutionized RA management, enabling targeted control of inflammation, deceleration of disease progression, and improved physical function (1–4). Furthermore, DMARDs have been shown to mitigate structural damage, alleviate symptoms, and improve the quality of life in patients with RA (1–4). Among DMARDs, methotrexate (MTX) remains the first line therapy (8), due to its proven efficacy, safety, and tolerability (1, 4, 9, 10). Its efficacy is hypothesized to be rooted in a multifaceted interplay of anti-inflammatory effects targeting various pathogenic cell types. Proposed mechanisms include its function as a folate antagonist, inhibition of cellular proliferation, enhancement of anti-inflammatory adenosine signaling, and modulation of critical inflammatory pathways such as the JAK/STAT pathway and NF-κB pathway (1, 11–13). Additionally, MTX helps restore CD4^+^ T lymphocyte balance, improving immune regulation in RA (6, 14–19). Yet, many patients fail to achieve an adequate therapeutic response (1, 13, 20, 21). Consequently, RA patients require monitoring for up to three months to assess their clinical outcomes and define the following treatment strategy. Therefore, identifying biomarkers, such as cellular response parameters, is essential for predicting treatment outcomes, optimizing therapeutic strategies, and ultimately improving patient health.

In summary, the efficacy of MTX in RA may result from a complex interplay of anti-inflammatory effects on various pathogenic cell types. To describe the immune cell changes occurring in vivo following MTX treatment we employed single-cell RNA sequencing (scRNA-seq) in conjunction with multi-color immunophenotyping of peripheral blood mononuclear cells (PBMCs) from treatment-naïve RA patients, compared to healthy controls (HC). Follow-up analyses were conducted at 3, 6, and 12 weeks after the initiation of MTX treatment. Using this time-resolved approach of cellular profiling, we uncovered changes in the abundances of distinct PBMC subsets, such as plasmablasts and T follicular helper (Tfh) cells throughout treatment, many of which closely resemble non-diseased phenotypes.

Importantly, our analysis revealed an early MTX-induced cellular response, preceding detectable clinical outcomes. The early responder cell types identified may serve as potential biomarkers for predicting therapeutic response to MTX. Finally, to benefit the broader scientific community, we developed a cell browser web platform to visualize the data presented in our study. Using the platform, expression profiles of specific genes can be investigated and visualized in a time-resolved manner following MTX treatment, facilitating further validation of the identified gene expression in the context of MTX treatment.

## Methods

### Patients and interventions

12 participants from the interdisciplinary outpatient clinic at the Division of Rheumatology at the Medical University of Vienna were enrolled in this study, comprising 6 HC and 6 RA patients diagnosed according to the American College of Rheumatology (ACR/EULAR) criteria for RA, as defined by classification criteria (22). All patients gave their written informed consent to retrospective data analysis as well as biobanking of blood samples. Samples were stored at the biobank at the Medical University of Vienna, a centralized facility for the preparation and storage of biomaterial with certified quality management (International Organization for Standardization (ISO) 9001:2015) (23). All participants were over 18 years of age. Ethical approval for this study was granted by the ethics committee of the Medical University of Vienna, Austria (1075/2021; 2071/2020). Patients or the public were not involved in the design, conduct, reporting, or dissemination plans of our research.

The RA group consisted of 2 male and 4 female patients, with a mean age of 55.83 ± 9.11 years and a mean Body Mass Index (BMI) of 27.62 ± 2.95. All of them were smokers or had a history of smoking (1 former smoker; ex-smoker for 5 years, and 5 current smokers). Patient characteristics, including laboratory markers, are summarized in Table 1. HC were matched for sex, age (52.5 ± 5.9), BMI (29.29 ± 4.2), and smoking status (1 former smoker, ex-smoker for 7 years, pack years: 14,33±11,5) to ensure comparability. In the HC cohort, an existing autoimmune disease was excluded based on the medical history. Additionally, HC did not exhibit positive autoantibodies (ANA, ENA, thyroid-associated antibodies) at the time of PBMC collection. All participants were of Caucasian descent.

**Table 1.**
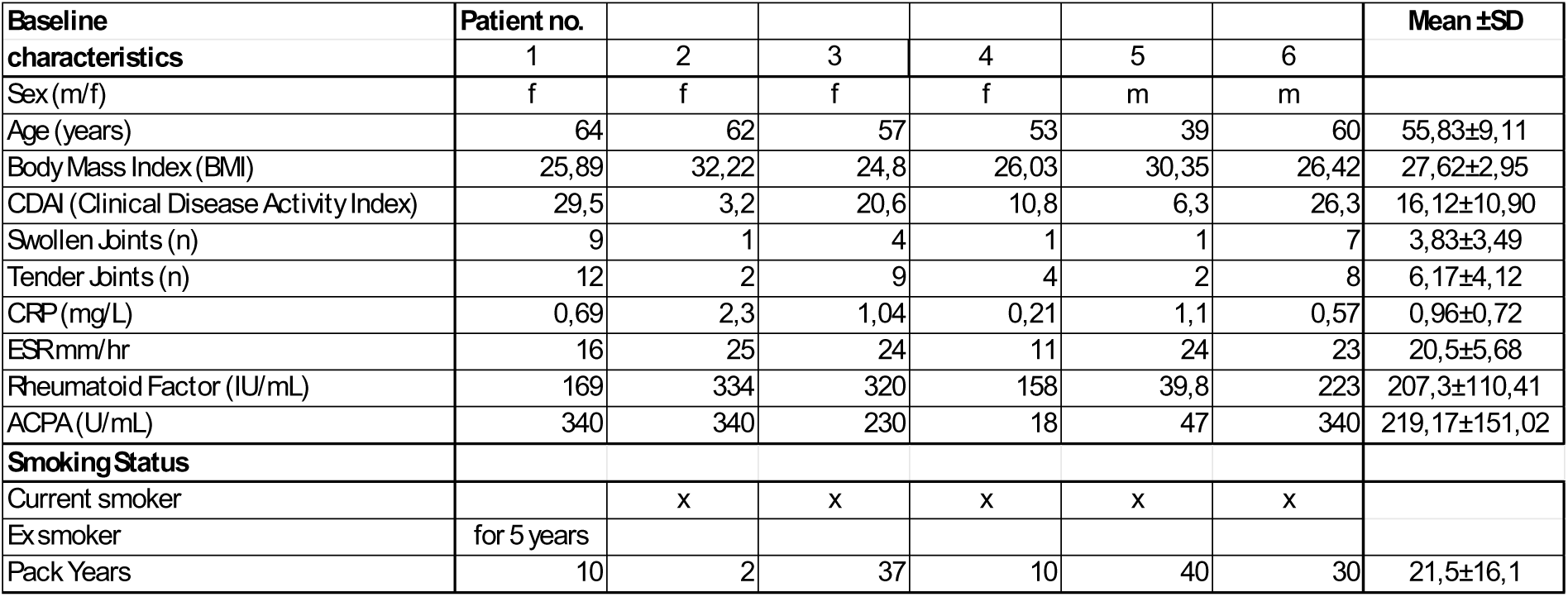
Baseline characteristics of RA patients.

A baseline blood collection was performed for all participants. Prior to the baseline blood collection (week 0), none of the participants had received corticosteroids in their lifetime. Additionally, all participants had used over-the-counter (OTC) nonsteroidal anti-inflammatory drugs (NSAIDs) for pain management.

For the RA patients, treatment with methotrexate (15 mg/week for the first 3 weeks, followed by a dose increase to 25 mg/week) was initiated. Three additional blood collections were performed at 3, 6, and 12 weeks following the initiation of treatment. Disease activity, as assessed by the Clinical Disease Activity Index (CDAI) at baseline and week 12 (24). All patients demonstrated a clear response to MTX, as delineated by Aletaha et al (8).

### Isolation and freezing of PBMCs

Blood was collected into heparin-containing tubes, and peripheral blood mononuclear cells (PBMCs) were isolated using Pancoll density gradient centrifugation. Briefly, 10 mL of whole blood was diluted at a 1:1 ratio with PBS (Sigma), layered on top of a 15 mL Pancoll (PAN-Biotech) solution, and centrifuged at 530 g for 22 minutes at room temperature (RT) without interruption. Following centrifugation, the distinct white layer containing PBMCs was carefully transferred into a new 50 mL Falcon tube, washed with PBS, and centrifuged at 400 g for 8 minutes at 4°C. The resulting cell pellets were resuspended in 5 mL of PBS. Human PBMCs were enumerated within a diameter range of 7 to 15 µm using a Z2 Coulter Particle Count and Size Analyzer (Beckman Coulter). Subsequently, PBMCs were pelleted (400 g, 8 min, 4°C), resuspended at a concentration of 10 - 20 x 10^6^/mL in freezing medium (RPMI-1640 supplemented with 20% FCS (Gibco) and 15% DMSO (Sigma)), transferred into cryovials, and stored in a CoolCell FTS30 cell freezing container for 24 hours. Afterward, the cryovials were transferred to liquid nitrogen for long-term storage.

### Thawing of PBMCs

10 x 10^6^ PBMCs dissolved in a freezing medium were incubated at 37°C for 5 minutes. Subsequently, the thawed contents were transferred to a 50 mL Falcon tube. Cell culture medium (RPMI-1640 supplemented with 10% FCS (GIBCO), 1% Penicillin/Streptomycin (GIBCO), and 1% GlutaMAX (GIBCO)) was added drop-wise until reaching a total volume of 50 mL, with a 1-minute incubation and mixing step following each duplication of the total volume. Following centrifugation (400 g, 5 minutes, 4°C) and removal of the supernatant, the resulting cell pellets were resuspended in 1 mL PBS and counted using a Coulter counter, following the previously described method.

### Sample processing for flow sorting and immunophenotyping

The PBMC samples from one HC and one time course of a RA patient (baseline, week 3, week 6, week 12) were consistently processed in parallel on the same day. Following the thawing of PBMCs and determination of cell numbers, 2 x 10^6^ PBMCs were set aside as unstained / Fluorescence-minus-one (FMO) controls for immunophenotyping. 4 x 10^6^ PBMCs were stained for viability using the L/D Fixable Blue Dead Cell Stain Kit (Thermo Fisher Scientific). In brief, a fresh aliquot of the L/D Fixable Blue Dead Cell Stain Kit was pre-diluted to a ratio of 1:40. PBMCs were diluted to a concentration of 1 x 10^6^ cells per 350 µL PBS, and the pre-diluted L/D dye was added at a ratio of 1:70. After vortexing, the sample was incubated in the dark for 15 minutes at room temperature. Subsequently, the samples were washed with 2% Buffer (PBS supplemented with 2% FCS) and resuspended in 250 µl of 2% Buffer. Following this, 300.000 viable PBMCs were sorted on a BD FACSAria™ Fusion Flow Cytometer, followed by a quality control step for purity using flow cytometry. The remaining unsorted PBMCs were then subjected to immunophenotyping.

### Preparation for 10x Genomics single-cell RNA sequencing

Freshly sorted PBMC samples from the time course of a single RA patient (baseline, week 3, week 6, week 12) were incubated with commercially available DNA-labeled antibodies (TotalSeq-A, A0251-A0254, BioLegend) at a concentration of 1 µL of antibody per 10 x 10^6^ PBMCs. Following a 30-minute incubation at 4°C, PBMCs underwent three washes with 2% Buffer. After the final wash, PBMCs were resuspended in 2% Buffer. The samples from these four time points were pooled for processing as a single sample per the manufacturer’s protocol, leveraging the antibody-linked barcodes for sample-specific demultiplexing of the sequencing data. Conversely, the corresponding HC sample was treated as an individual sample for scRNA-seq. Libraries for scRNA-seq were generated using the Chromium Controller and the Next GEM Single Cell 3’ Reagent Kit (v3 or v3.1, 10x Genomics) following the manufacturer’s instructions. The libraries were then sequenced by the Biomedical Sequencing Facility (BSF) at the Research Center for Molecular Medicine of the Austrian Academy of Sciences (CeMM) using the Illumina HiSeq 3000/4000 platform.

### Single-cell RNA sequencing analysis

Raw sequencing data underwent pre-processing and alignment to the GRCh38 human reference genome as well as demultiplexing of the antibody-linked barcodes using Cell Ranger (v6.1.2, 10x Genomics) generating count data. Cells with fewer than 300 features and more than 10% mitochondrial counts, as well as cells with ambiguous sets of markers (for example, both CD19 and CD3) were filtered out of the analysis, resulting in 65269 cells used for downstream analysis. We used the Seurat v5.0.01 package (25) for data normalization (using log normalization with a scale factor of 10000) and scaling and constructed a PCA based on 2,000 most variable features. A Shared Nearest Neighbor (SNN) graph was constructed based on the first 14 PC dimensions with euclidean distance as a distance dimension. Clusters were obtained based on the previously constructed SNN graph using a Louvain algorithm and a resolution of 0.5. A UMAP was constructed using Seurat’s default parameters and based on the first 14 dimensions. Per-cluster marker genes were found using the FindAllMarkers function from Seurat. Clusters were annotated based on the expression of top marker genes and an automated annotation pipeline using SingleR (25) with the Monaco dataset as a reference (26). To obtain the cluster subsets and substates, the Seurat object was subsetted into three categories: T cells, B cells, and Myeloid cells. As the batch effect was strongly driving the variance in the subsetted data, each of the subsets was integrated by batch using an anchor-based CCA integration. The subsets were annotated using the strategy described earlier. The top marker genes for each sub-cluster are denoted in Supplementary Figure 3 and a full list is provided in Supplementary Table 2.

The proportional difference in cell populations was calculated pairwise for each sample again the corresponding RA baseline sample using the scProportionTest package in R, using permutation with replacement for significance testing as outlined in [https://github.com/rpolicastro/scProportionTest/releases/tag/v1.0.0].

Differential expression analysis was conducted in a pairwise manner using the FindMarkers function in Seurat, using a logistic regression method “LR” and the batch effect (corresponding to donor ID) as a latent variable. Genes were deemed significant if the BH-corrected FDR was below 0.05 and the absolute logarithm of the fold change was greater than 0.1. The analysis was conducted using both the main cell types and the subsets as labels and results were aggregated by cell type and condition.

### Enrichment analysis and enrichment term clustering

Differentially expressed genes for either HC against baseline RA patients or MTX treatment time-course were aggregated by cell type. The results were then functionally enriched using the enrichR package in R and clustered and summarized using the SummArIzeR tool we have developed (27). Briefly, the databases used for enrichment analyses were: “BioPlanet_2019”, “GO_Biological_Process_2023”, “Reactome_2022”, “MSigDB_Hallmark_2020” and “DisGeNET” from enrichR. Top 10 terms per database and comparison were selected and terms with fewer than 5 genes were filtered out. The term similarity scores were then calculated based on Jaccard similarity of genes and clusters were inferred from the similarity matrix using a Walktrap community finding algorithm, Edges with a threshold below 0.16 for the HC vs RA comparison and 0.24 for the MTX timecourse comparison were removed from the analysis. The term dendrogram and corresponding clusters are shown in Supplementary Figure 4.

### Gene Regulatory Network (GRN) construction from Single-cell RNA

To create gene regulatory networks and the differences between them we used the SCORPION package as described by Osorio et al (28). Briefly, all significantly different genes across any comparison from the previous steps were used to reduce the size of the datasets. The datasets were further split by patient and cell type and monocytes and CD4 memory T cells were selected for further analysis. The scorpion algorithm was then run for each sample using default parameters (gammaValue = 10, nPC = 25, assocMethod = “pearson”, alphaValue = 0.1, hammingValue = 0.001, nIter = Inf, zScaling = TRUE, randomizationMethod = “None”). Interaction weights were summarized by group and network summary, comparison, statistical analysis, and plotting were performed using methods and functions specified in (29). A p-value below 0.05 was considered for filtering meaningful interactions. Groups were compared pairwise and only interaction pairs where at least one pairwise comparison was statistically significant were retained. Interaction pairs were further filtered for readability, such as at least one of the interactors is significantly differentially perturbed in the corresponding cell type. The hub genes were defined as genes with the highest level of centrality for a corresponding network.

### PBMC immunophenotyping staining

Staining of PBMCs was performed as described in a previously published study (30). The clones, manufacturers, and catalog numbers of the antibodies used are detailed below.

### Import of immunophenotyping data and clean-up

The immunophenotyping dataset was imported into FlowJo™ software (version 10) in .fcs file format. Initially, all samples underwent data cleanup using the FlowAI plugin within FlowJo™. FlowAI was applied to all uncompensated parameters, with the following settings: Anomalies to exclude were defined as Flow rate & dynamic range; Second fraction FR was set to 0.100; Alpha FR was set to 0.0100; Maximum changepoints were limited to 3; Changepoint penalty was set at 200; Dynamic range check side was set to Both. Additionally, outliers were automatically removed by the software. Notably, less than 5% of events were excluded from all samples through the application of FlowAI. The events identified as “good events” by the FlowAI software were utilized for subsequent analyses.

### Analysis of the immunophenotyping dataset in FlowJo™

After successful data cleanup, all samples underwent manual pre-gating in FlowJo™ to eliminate residual aggregates, debris, and doublets, based on evaluation of the scatter profiles. Subsequently, manual gating for distinct peripheral blood mononuclear cell (PBMC) populations was performed, guided by relevant literature references (30–34). Dimensionality reduction was then implemented using the UMAP plugin within FlowJo. Initially, to visualize major PBMC populations (Figure 1B), UMAP analysis was conducted on viable cells, employing a panel of compensated fluorescent parameters including CD16, CCR5, CCR4, CD11c, CD56, CD8, CCR7, CD123, CD161, IgD, CD20, CD3, IgM, CD28, CCR6, CXCR5, PD-1, CD141, CD57, CD14, CD11b, TCRγδ, CCR10, CD4, CD24, OX40, CXCR3, CD25, CD27, CD1c, CD19, CD127, HLA-DR, and CD38. Additionally, to visualize B cell subsets, UMAP analysis was performed on CD19^+^ CD20^+/-^ B cells using specific compensated fluorescent parameters: IgD, CD20, CD27, IgM, CXCR5, PD-1, OX40, Fas, CD25, HLA-DR, and CD38. For visualization of T cell subtypes, UMAP analysis was conducted on CD3^+^ T cells, employing a panel of compensated fluorescent parameters including CCR5, CCR4, CD11c, CD56, CD8, CCR7, CD123, CD161, CD28, CCR6, CCR4, CXCR5, CXCR3, PD-1, CD141, CD57, CD11b, TCRγδ, CCR10, CD4, CD24, OX40, Fas, CD25, CD27, CD1c, CD127, HLA-DR, and CD38. To delineate different myeloid subsets, UMAP analysis was performed on CD45^+^ CD3^-^ CD19^-^ viable cells, utilizing the compensated fluorescent parameters CD16, CD11c, CD123, CD141, CD14, CD11b, CD1c, HLA-DR, and CD38. Subsequently, percentages of the respective populations were extracted and exported in .csv format for further downstream analysis.

**Figure 1.**
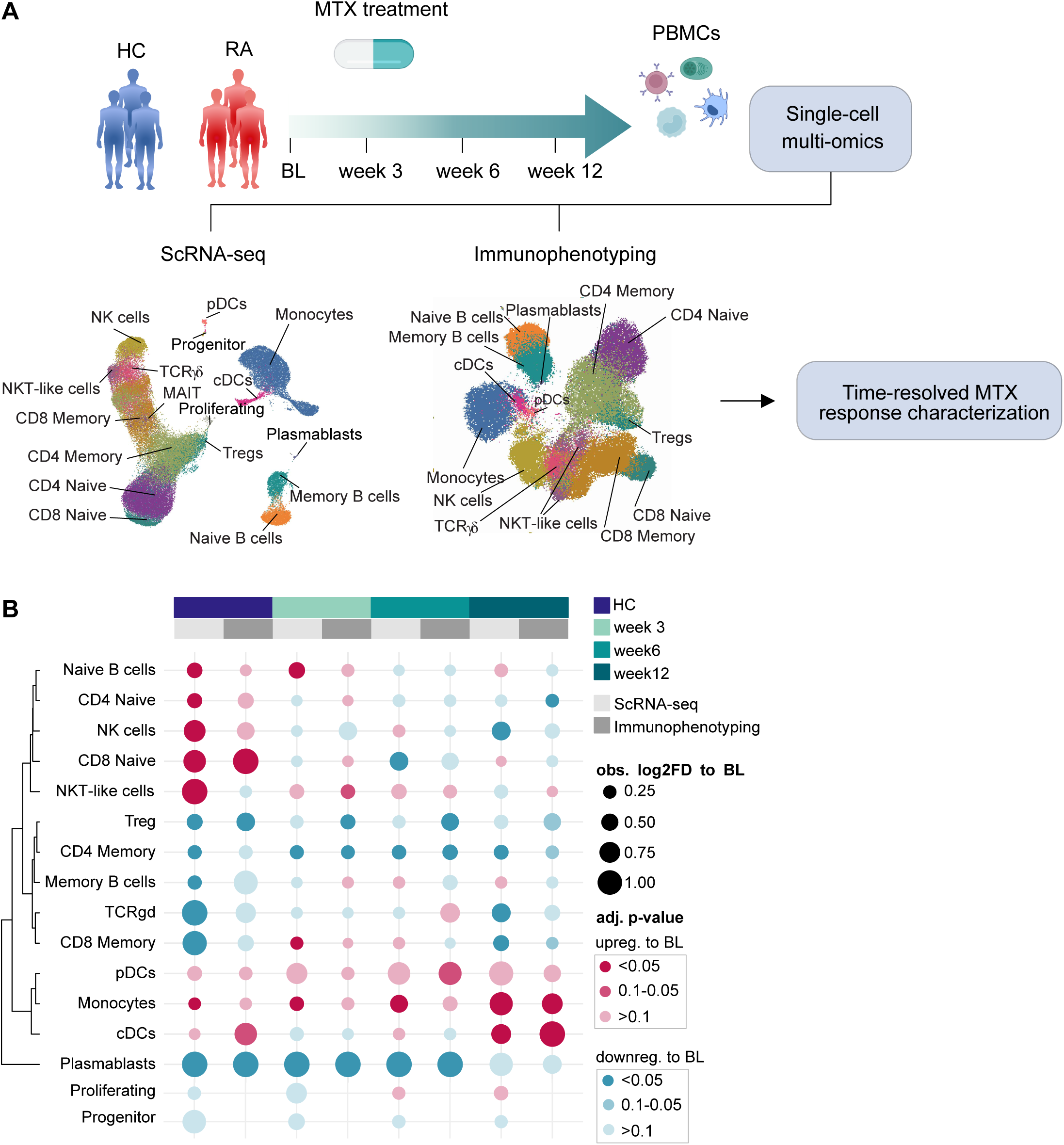
Multimodal analysis of PBMC subset perturbation during Methotrexate therapy in RA patients. (A) Schematic representation of the study design and sampling timepoints. PBMCs have been isolated from six treatment-naive patients (baseline), and after 3, 6 and 12 weeks from the initiation of MTX therapy, as well as six age- and sex-matched controls. The cells were subjected to immunophenotyping (per group n = 6) and single-cell RNA sequencing (per group n = 5). Uniform Manifold Approximation and Projection (UMAP) plots are shown representing the global cell types detected using the two technologies. (B) Cell type abundance comparison across the two methodologies and cell types. HC and each MTX treatment timepoint (3, 6 and 12 weeks post MTX) were compared to baseline RA (BL). Cell types with an increase in abundance compared to BL are depicted in red, whereas decrease in blue. The size of the dots corresponds to the absolute observed log2 fold difference (log2FD) compared to BL, whereas the color intensity represents the adjusted p-value using a permutation test. The log2FD values are capped at 1. Abbreviations: healthy controls (HC), rheumatoid arthritis (RA), baseline (BL), Natural Killer T cell (NKT) – like cells, Methotrexate (MTX), gamma-delta T cell (TCRγδ), conventional Dendritic Cells (cDC), plasmacytoid Dendritic Cells (pDC).

### Downstream data analysis and statistics in R

The subsequent data analysis was conducted in RStudio (R Development Core Team). Data from .csv files were imported into the R environment using standard import functions. Boxplots, bar charts, and line plots were generated using the “ggplot2” and “plotly” R packages. Statistical significance was assessed in RStudio using the “rstatix” R package. For paired comparisons during the time course of MTX treatment in RA patients, paired t-tests were employed, along with Benjamini-Hochberg adjustment and a 90% confidence interval. Significance was defined as follows: ^#^P<0.1, *P<0.05, **P<0.01, ***P<0.001, and ****P<0.0001. To compare HC and RA patients, an unpaired t-test was utilized, with significance determined using the same criteria.

### Reagents

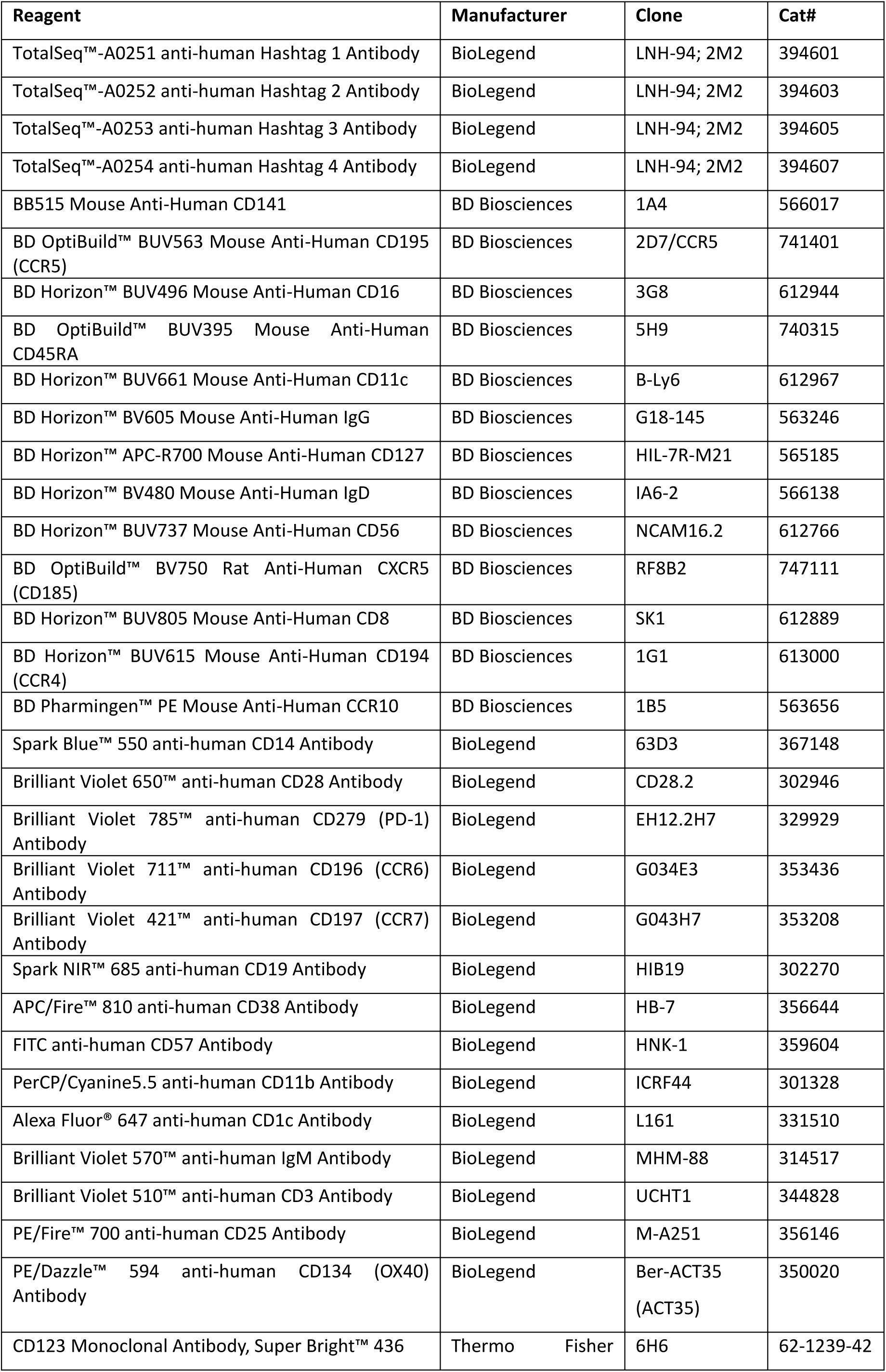
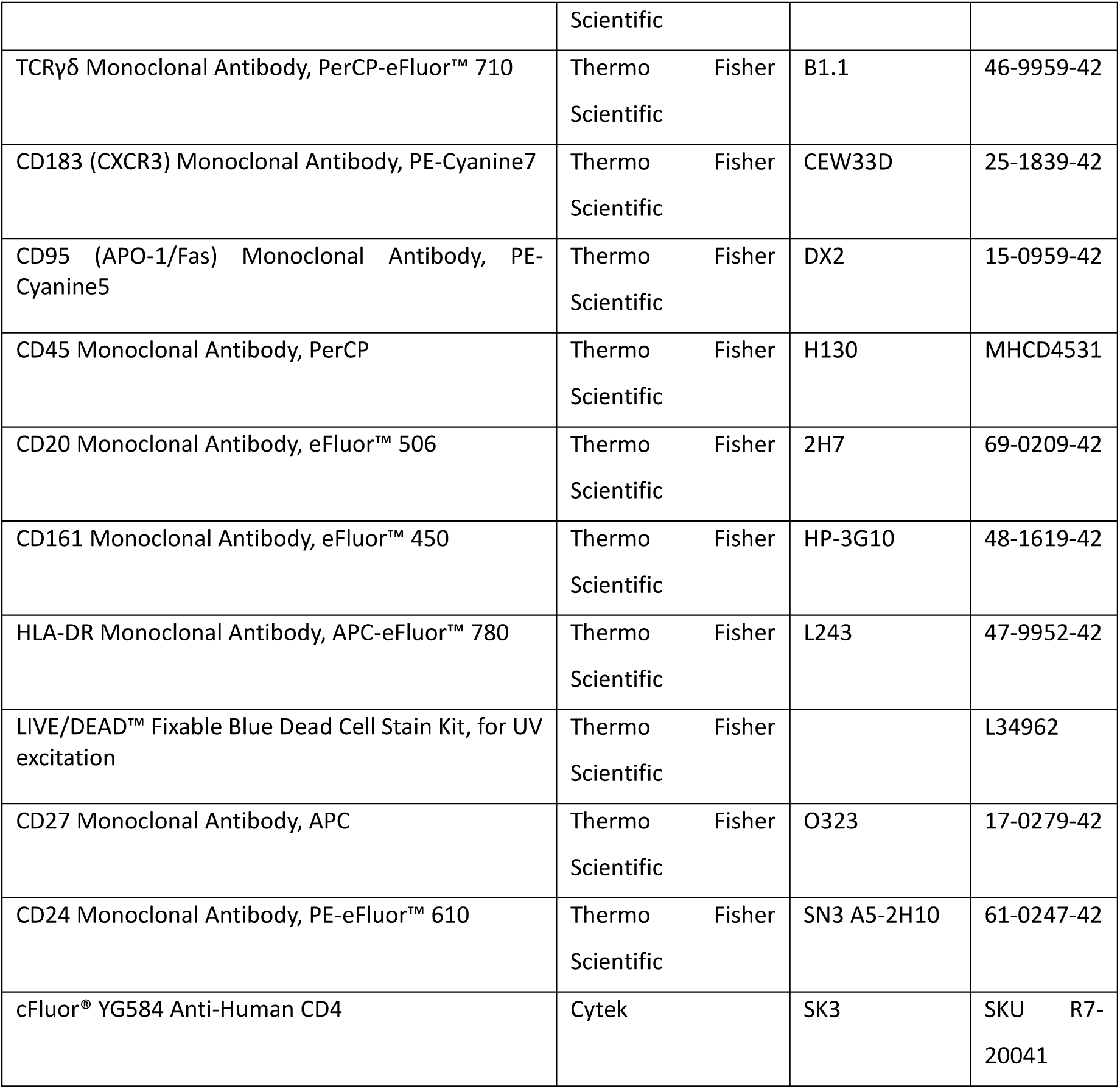

## Results

### Multi-omics approach reveals perturbed PBMC subsets in patients with RA as a response to methotrexate treatment

In the presented experimental setup, we recruited five treatment-naive RA patients (three female, two male) and five healthy controls. The patients’ demographic details and clinical/laboratory data are summarized in Table **1**. To investigate cellular and transcriptional alterations in the immunological landscape of RA patients induced by MTX treatment, we collected PBMCs from HC and RA patients before initiation of therapy (baseline, BL), as well as 3, 6, and 12 weeks after the start of MTX treatment **(Figure 1A)**. PBMCs were analyzed by spectral cytometry for immunophenotyping. Sorted viable cells were subjected to scRNA-seq **(Figure 1A)**. **Supplementary figures 1 and 2** illustrate the gating strategy used for immunophenotyping and the effects of various variables on the cell-type clustering for scRNA-seq, respectively.

The two complementary methodological approaches, immunophenotyping and scRNA-seq **(Figure 1B)**, allowed for the identification of distinct cell populations representing major PBMC subtypes, including naïve and memory CD4^+^ T cells, naïve and memory CD8^+^ T cells, TCRγδ T cells, monocytes, naive and memory B cells, as well as plasmablasts, conventional dendritic cell (cDC), plasmacytoid dendritic cells (pDC), natural killer (NK) cells, and natural killer T-like (NKT-like) cells. Notably, scRNA-seq analysis identified two additional cell types—proliferating cells and progenitor cells—when compared to spectral cytometry, as highlighted in **Figure 1B**.

To evaluate the longitudinal effects of MTX on cell type abundances and to compare the two methodologies a permutation test was applied, comparing each treatment timepoint to baseline. Moreover, baseline differences between HC and RA patients were calculated. Changes were represented as a log2 fold change (log2FC) to baseline **(Figure 1B)**. Circle sizes correspond to the absolute log2FC, with increased abundance represented in red and decreased abundance in blue. The intensity of the colors reflects the corresponding significance levels. Hence, we demonstrated the comparability between scRNA-seq and immunophenotyping in detecting PBMC populations, as the cellular abundances of different cell types exhibited a strong correlation between the two methods.

Hierarchical clustering enables a comparative analysis between RA patients and HC, while also highlighting trends across the longitudinal timeline. In this context, naïve PBMC subtypes, NK cells, and NKT-like cells were grouped into a “naïve” cluster, while memory cells (CD4^+^ memory T cells, CD8^+^ memory T cells, memory B cells), as well as tregs and TCRγδ T cells, formed a “memory” cluster. A third “myeloid” cluster contained DCs and monocytes, and the fourth cluster comprised plasmablasts.

Cells within the naïve cluster were upregulated in HC compared to RA baseline. Notably, by week 3 post-MTX treatment, the abundance of naïve B cells and NKT-like cells was increased when compared to RA baseline, reflecting proportions indicative of a healthy phenotype. In contrast, PBMC subsets within the memory cluster were less abundant in HC as compared to RA patients. Similar to the naïve cluster, memory PBMC subsets showed a reduction as early as 3 weeks post-MTX treatment, with these lower levels persisting throughout the study period. In the myeloid cluster, pDCs, cDCs, and monocytes were elevated in HC when compared to RA baseline. Following MTX treatment, these populations were also increased in RA patients, reaching levels comparable to HC. The most pronounced differences were observed in plasmablasts, which were significantly reduced in HC compared to RA patients. MTX therapy notably decreased this subset to healthy levels as early as 3 weeks post-treatment initiation.

In summary, these results provided a comparative analysis of cellular abundances between HC and RA patients, revealing that the PBMC landscape in RA was characterized by reduced levels of naïve cells and increased levels of memory phenotype cells.

### Lymphocyte and myeloid modulation by MTX in RA

We further employed a fine-grained subclustering of major PBMC subtypes to investigate MTX treatment’s potential impact on specific subpopulations. Distinct B cell subpopulations, including naïve B cells, non-switched memory B cells (NonSwMe Bc), switched-memory B cells (SwMe Bc), and plasmablasts **(Figure 2A)** could be identified using both immunophenotyping and scRNA-seq. Notably, naïve B cell frequencies were reduced in RA patients compared to HC, whereas switched-memory B cells and plasmablasts were markedly elevated **(Figure 2B)**. MTX treatment exerted profound effects on plasmablast proportions, with the most significant reduction observed as early as three weeks after treatment initiation **(Figure 2C)**. Further subsetting identified multiple cellular substates of naïve and non-switched memory B cells **(Supplementary Figure 3A)**, however, none of these subsets were markedly affected by MTX treatment. Immunophenotyping further facilitated the evaluation of B cell activation states using key markers such as CD25, CD38, and Fas **(Figure 2D)**. In untreated RA patients, B cells exhibited an elevated activation profile compared to HCs **(Figure 2D)**. Following MTX therapy, these activation markers demonstrated a progressive decline, culminating in an activation profile at week 12 that closely resembled that of HC **(Figure 2D)**.

**Figure 2.**
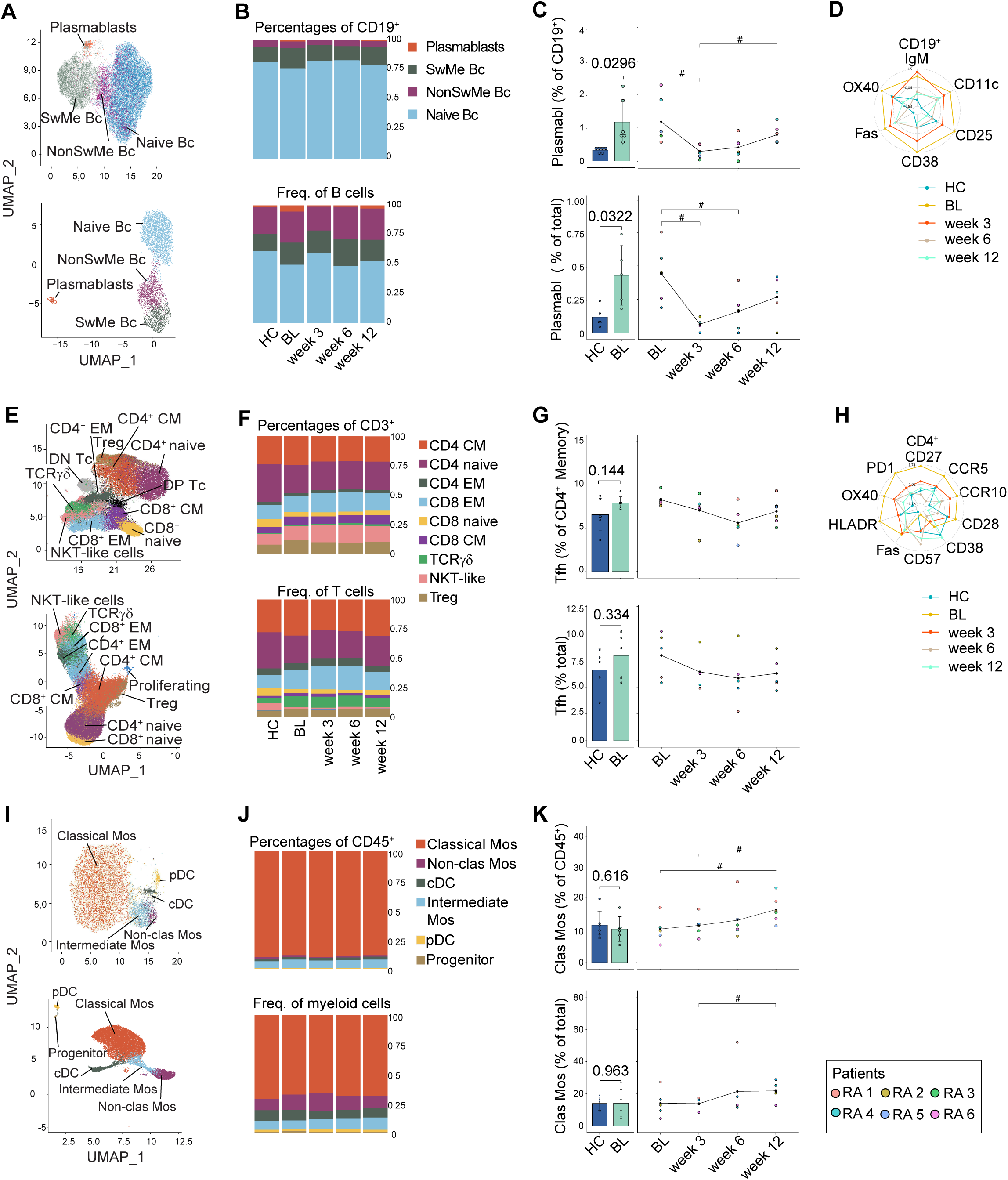
Dynamic modulation of immune cell subsets and activation markers during Methotrexate therapy in RA patients. (A, E, I) Uniform Manifold Approximation and Projection (UMAP) of (A) B cell subsets, (E) T cell subsets, and (I) myeloid cell populations from concatenated samples, HC and RA patients at baseline and during therapy (weeks 3, 6, and 12), illustrating subset distributions. UMAPs were generated using (upper panels) immunophenotyping and (lower panels) scRNA-seq data. (B, F, J) Frequencies of (B) B cell subpopulations (naive B cells, switched and non-switched memory B cells, plasmablasts), (F) T cell subpopulations (CD4^+^ central memory, CD4^+^ naive, CD4^+^ effector memory, CD8^+^ effector memory, CD8^+^ naive, CD8^+^ central memory, TCRγδ, NKT-like cells, regulatory T cells), and (J) myeloid subpopulations (classical monocytes, non-classical monocytes, intermediate monocytes, conventional dendritic cells, plasmacytoid dendritic cells, progenitors), expressed as a percentage of (upper panels) CD19^+^ cells (B), CD3^+^ cells (F), or CD45^+^ cells (J), and as a percentage of total cells (lower panels) across HC and RA patients at baseline and during therapy. (C, G, K) Bar charts and dot plots depict changes in (C) plasmablasts, (G) T follicular helper cells, and (K) classical monocytes, expressed as a percentage of (upper panels) CD19^+^ cells (C), CD4^+^ memory T cells (G), or CD45^+^ cells (K), and as a percentage of total cells (lower panels), respectively, between HC and RA patients throughout therapy. Data were generated using (upper panels) immunophenotyping and (lower panels) scRNA-seq. (D, H) Radar plots display longitudinal changes in activation markers for (D) B cells and (H) CD4^+^ T cells in HC and RA patients during therapy, generated by immunophenotyping. For paired comparisons during the MTX treatment time course in RA patients, paired t-tests with Benjamini-Hochberg adjustment were applied, using a 90% confidence interval. Significance thresholds were defined as follows: #P < 0.1, *P < 0.05, **P < 0.01, ***P < 0.001, ****P < 0.0001. Comparisons between HC and RA patients were conducted using unpaired t-tests, with the same significance criteria. Detailed statistical analyses and sample sizes are available in the Methods section. Abbreviations: healthy controls (HC), rheumatoid arthritis (RA), plasmablasts (Plasmabl), switched memory B cells (SwMe Bc), non-switched memory B cells (NonSwMe Bc), Central memory (CM), effector memory (EM), natural killer T cell (NKT) – like cells, T follicular helper (Tfh) cell, methotrexate (MTX), monocytes (Mos), classical monocytes (Clas Mos), non-classical monocytes (Non-clas Mos), conventional dendritic cells (cDC), plasmacytoid Dendritic Cells (pDC).

Using both immunophenotyping and scRNA-seq distinct subsets of CD3^+^ T cells were delineated and quantified. These subsets include CD4^+^ and CD8^+^ T cells with naïve phenotype, memory (CM) phenotype, or effector (EM) phenotype, as well as Tregs, double-positive (DP) T cells, double-negative (DN) T cells, TCRγδ T cells, and NKT-like cells **(Figure 2E)**. At baseline, RA patients exhibited a reduction in naïve CD4^+^ and CD8^+^ T cells, while central memory (CM) CD4^+^ and CD8^+^ T cells, Tregs, and NKT-like cells were elevated **(Figure 2F)**. MTX treatment restored T cell subsets to levels comparable to those in HC, evidenced by an increase in naïve T cells and a reduction in memory T cells **(Figure 2F)**. Moreover, MTX reduced the elevated levels of T follicular helper (Tfh) cells observed in RA patients, with the most substantial reductions occurring within the first three weeks of treatment **(Figure 2G)**. Further subsetting of T cells based on scRNA-seq information **(Supplementary Figure 3B)** has revealed differences in the proportion of multiple cell types between HC and RA at baseline. In contrast, only a few subsets (CD8 effector cells, MAIT, memory Tregs) exhibited significant changes during/after MTX treatment **(Supplementary Figure 3D)**. To explore the therapeutic potential of MTX beyond its impact on cell abundances, we assessed T cell activation states using protein expression data. This analysis demonstrated elevated levels of key activation markers, including Fas, HLA-DR, OX40, and PD-1, on CD4^+^ T cells in RA patients compared to HC **(Figure 2H)**. By week 3 of MTX therapy, activation markers were significantly reduced, resulting in an activation profile resembling that of HC. This alignment was sustained through week 12, underscoring the multifaceted therapeutic mechanisms of MTX. **(Figure 2H)**.

A fine-grained analysis of the myeloid subset combining both immunophenotyping and scRNA-seq confirmed established myeloid subset distributions, such as classical monocytes, intermediate monocytes, and non-classical monocytes, as well as cDCs, pDCs, and progenitors (CD34^+^ hematopoietic progenitor cells) (26) **(Figure 2I)**. Classical monocytes and cDCs exhibited a slight reduction in RA patients compared to HC **(Figure 2J**, **2K)**, and following MTX treatment, the proportions of these subsets increased progressively over time **(Figure 2J**, **2K)**. Further classification of the classical monocyte population was performed on a scRNA-seq level **(Supplementary Figure 3C)** A significant increase, induced by MTX treatment was demonstrated in the IL1B subset after 3 and 6 weeks of treatment, as well as in the IFN subset after 12 weeks of treatment **(Supplementary Figure 3D)**. The IL1B subset was significantly increased in PBMCs from HC compared to RA patients **(Supplementary Figure 3D)**.

In brief, MTX treatment profoundly modulated immune cell dynamics, progressively normalizing lymphocyte, and myeloid subsets to levels comparable to HC. These effects included rapid reductions in activation markers and shifts in cellular proportions observed as early as three weeks post-treatment initiation. This underscores the multifaceted mechanisms of MTX treatment, potentially occurring well before the onset of a clinical response.

### MTX targets multiple pathways and gene hubs over time

We next conducted a differential expression analysis comparing cell-type specific perturbations across the MTX treatment time course as well as the baseline transcriptional perturbations in RA compared to HC. A full list of perturbed genes is shown in **supplementary table 2**. Genes were selected if the pairwise comparison p-value was below 0.05 and an absolute log2-fold difference of at least 0.1 was detected. Enrichment analysis was performed to summarize the expressed genes into relevant functional categories **(Figure 3A)**. A comparison of HCs and RA patients confirmed that modalities related to autoimmune diseases and inflammation were perturbed across multiple cell types. Notably, pathways related to translation, ribosome function, and ribosome biogenesis were broadly altered across all PBMC subsets, whereas pathways such as IFN-α and splicing and cell cycle modulation were specifically affected in memory and naive CD4^+^ T cells. Monocytes were particularly enriched in autoimmune and antigen processing pathways, TNF-α and juvenile autoimmune diseases, IFN-γ response, and other inflammatory responses **(Figure 3A)**. A detailed breakdown of the terms within each enrichment category is provided in **Supplementary Figure 4A**.

**Figure 3.**
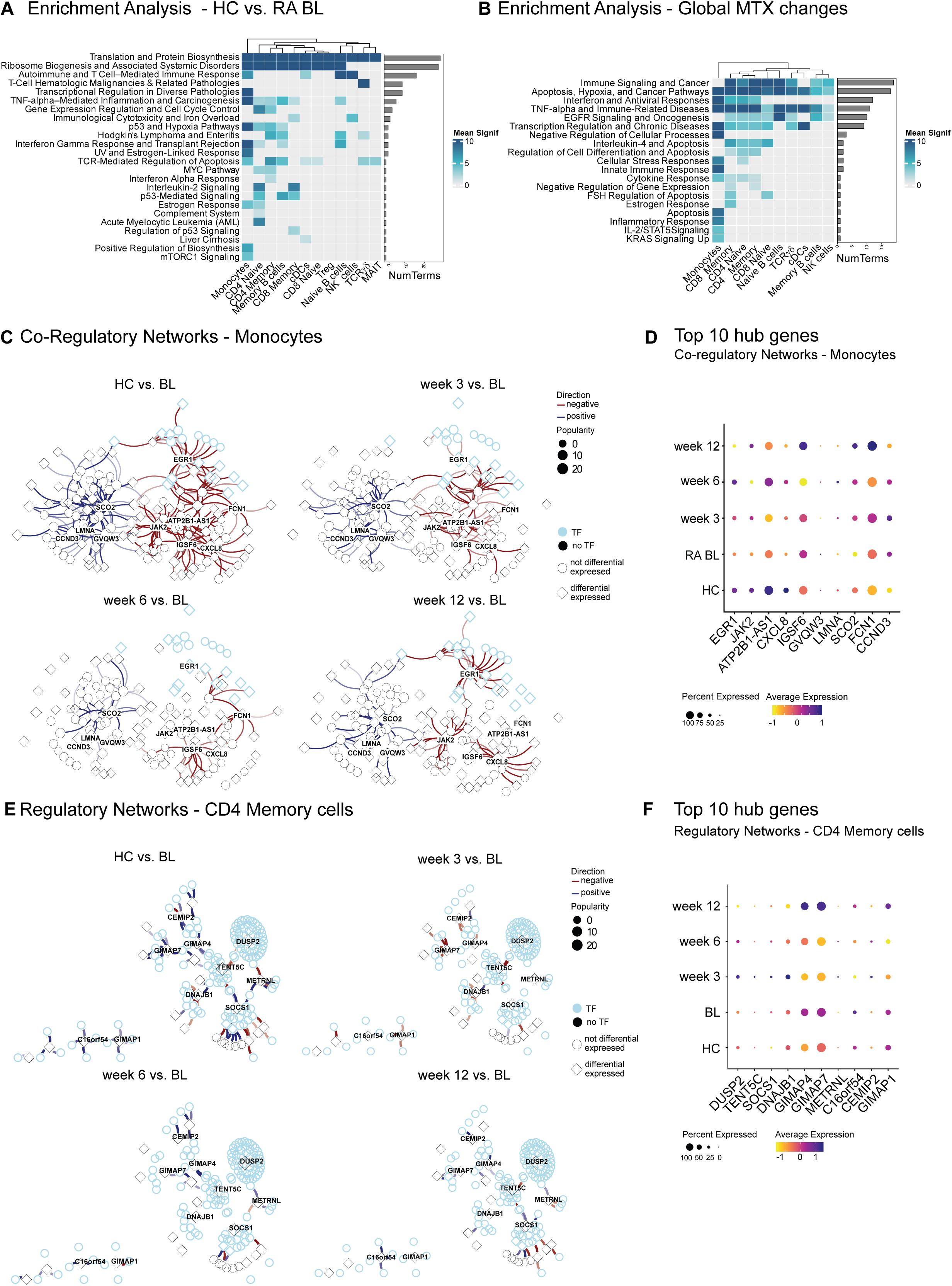
Pathway and gene regulation perturbation by Methotrexate in PBMC subsets. (A and B) Summarized enrichment results for cell-type specific differentially expressed genes. Five databases were used for the enrichment: “BioPlanet_2019”, “GO_Biological_Process_2023”,“Reactome_2022”, “MSigDB_Hallmark_2020”, “DisGeNET”, and the top 10 most significant terms were selected. Genes were considered differentially expressed if they reached an absolute log2 fold change of 0.1 and adjusted p-value below 0.05. Color represents the negative log10 of the averaged adjusted p-value per cluster. (A) Summarized enrichment for differentially expressed genes between HC and RA BL. (B) Summarized enrichment for genes affected by MTX treatment. (C) Co-regulatory network in monocytes; edges depict the difference in co-expression weights between either HC and RA BL or weeks 3, 6, and 12 of MTX treatment and RA BL, with red being a weaker relationship and blue stronger. The intensity of the color represents the magnitude of the difference. Light blue nodes represent transcription factors and diamond-shaped nodes represent genes affected by MTX treatment in the corresponding cell type. Hub genes are highlighted. (D) Hub genes corresponding to figure 3C (supplementary figure 5B). Color depicts the scaled average expression of the gene in monocytes across groups. Dot size represents the percentage of cells with non-zero expression of the gene. (E) Regulatory network in CD4 memory T cells; edges depict difference in co-regulation weights of two genes, one of which is a transcription factor, between either HC and RA BL or weeks 3, 6, and 12 of MTX treatment and RA BL, with red being a weaker relationship and blue stronger. The intensity of the color represents the magnitude of the difference. Light blue nodes represent transcription factors and diamond-shaped nodes represent genes affected by MTX treatment in the corresponding cell type. Hub genes are highlighted. (F) Hub genes corresponding to figure 3E (supplementary figure 6A). Color depicts the scaled average expression of the gene in CD4 memory T cells across groups. Dot size represents the percentage of cells with non-zero expression of the gene.

MTX treatment modulated immune signaling and cancer pathways, as well as apoptosis pathways, hypoxia pathways, and cancer pathways across multiple cell types. The affected genes were associated with pathways such as TGF-β, IL1, IL2, and TLR signaling **(Figure 3B and Supplementary Figure 4B)**, suggesting a potential anti-inflammatory mode of action. Furthermore, MTX therapy broadly influenced genes linked to TNF signaling and autoimmune diseases **(Figure 3B)**, with RA emerging as one of the key enriched terms within this cluster **(Supplementary Figure 4B)**. Cytokine and innate immune response, along with interferon and antiviral responses and proliferation, were specifically affected in monocytes. In contrast, IL4 signaling, apoptosis, and regulation of cell differentiation were predominantly impacted in naive and memory T cells.

We further aimed to identify the key players and gene regulatory changes associated with RA in comparison to a healthy phenotype. Additionally, we sought to investigate the time-resolved effects of MTX treatment on these regulatory alterations. We focused on two main cell types—monocytes and CD4^+^ memory T cells—since these cells exhibited significant changes in both cell proportions **(Figure 1**, **Figure 2)** and gene expression following MTX treatment **(Figure 3B)**.

**Figure 3C** illustrates a simplified gene co-regulatory network (GcRN) in monocytes, with embedded nodes representing co-expressed genes (the full network is shown in **Supplementary Figure 5B**). Transcription factors (TF) are represented by light-blue nodes, while diamond-shaped nodes indicate MTX-perturbed genes in monocytes. Highlighted genes were identified as hub genes based on their highest connectivity. The upper left panel (HC vs BL) depicts the differences in co-expression intensity between gene pairs in HC when compared to RA baseline, with red edges representing lower intensity and dark blue higher intensity. To evaluate the impact of MTX treatment on the dynamic alteration of the gene regulatory network we compared MTX-treated RA samples to baseline RA samples in a time-resolved manner (at 3, 6, and 12 weeks after MTX treatment). The most pronounced regulatory differences were detected between HC and RA BL. MTX treatment corrected the gene regulatory disturbances in RA, restoring a network profile resembling that of a healthy phenotype. The expression dynamics of the identified regulatory hub genes are shown in **Figure 3D**. Notably, four of the ten hub genes (EGR1, JAK2, CXCL8, FCN1) have previously been implicated in RA pathogenesis (35–37).

A gene regulatory network (GRN) of TF - target interaction in monocytes revealed a more complex interaction pattern **(Supplementary Figure 5A)**, indicating that MTX modulates the expression of immune response-related genes such as TNFAIP3 and NFKBIZ **(Supplementary Figure 5A, Supplementary Figure 7A)**. CD4^+^ Memory T cells, in contrast, exhibit a centralized co-regulatory network **(Supplementary Figure 6B)** with SOCS1 identified as a key regulatory hub gene. In addition, SOCS1, MNDA, DUSP5, and DUSP1 **(Supplementary Figure 7B)** are known to exert strong immunomodulatory effects and may play a potential role in RA pathogenesis (38–41).

**Figure 3E** represents a simplified GRN in CD4^+^ memory T cells, with embedded nodes representing co-regulated genes (the full network is shown in **Supplementary Figure 6B**). The GRN in CD4^+^ memory cells displays modular regulation with DUSP2, SOCS1, GIMAP4, GIMAP7, and GIMAP1 acting as central hubs **(Figure 3E, F, Supplementary Figure 6A)**. The gene regulatory network exhibited a similar trend in dynamics to that observed in monocytes (**Figure 3E)**.

Taken together, these data suggest that transcriptional perturbations in immune cells induced by RA can be restored by MTX treatment.

## Discussion

RA is a multifactorial autoimmune disease characterized by chronic joint inflammation, leading to cartilage destruction and bone erosion (1–4). Disease-modifying anti-rheumatic drugs (DMARDs) have transformed RA management by targeting inflammation, reducing structural progression, and improving physical function (1–4). MTX remains the cornerstone of RA treatment due to its excellent efficacy, safety, and toxicity profile. Its known mechanisms of action include anti-inflammatory, anti-metabolic, and anti-proliferative effects (1, 4, 9, 10). Despite significant advancements in understanding the effects of MTX, the precise molecular and cellular mechanisms underlying its therapeutic efficacy in RA remain incompletely elucidated. In this study, we sought to unravel these mechanisms by employing an integrative multi-omics approach, incorporating single-cell RNA sequencing (scRNA-seq) and comprehensive immunophenotyping. Through this novel methodological framework, we aimed to expand the existing knowledge and generate new insights into the therapeutic actions of MTX in RA.

Our findings show that MTX reshapes immune cell populations and gene expression profiles in RA, particularly affecting T cells and monocytes. The resulting phenotype closely resembles that found in healthy individuals.

Differential gene expression analysis highlighted key transcriptional changes associated with RA and MTX treatment spanning multiple immune cell subsets. RA was characterized by significant alterations in pathways related to autoimmune diseases, inflammation, ribosomal function, and cytokine signaling compared to healthy phenotype. These findings align with existing literature highlighting the central role of chronic immune activation in RA pathogenesis (42, 43). Notably, we could demonstrate that MTX treatment targets RA-specific pathways across a broad range of immune cell types. So far, only a small number of studies have explored the transcriptomic changes induced by MTX on PBMCs (20), however, the findings are limited to bulk PBMC samples and usually only include a later sampling time point of 12 weeks and beyond. One other study has so far performed scRNA-seq, similarly limited to a single late time point (44). Our study, in contrast, shows early-onset changes in the immune cell landscape induced by MTX and robust transcriptional perturbations as early as 3 weeks following treatment initiation. This indicates that cellular and molecular effects of MTX may arise before any observable clinical improvement. In addition, a recent study showed that active MTX metabolites such as MTX-PG1 and MTX-PG2 could be detected in PBMCs as early as one week after initiation of MTX treatment (45).

Building on these early changes, we compared the cellular landscape before initiating MTX treatment. We identified elevated numbers of plasmablasts in the RA baseline compared to HC, while a reduction was detected in week 3 after MTX treatment. Further, MTX modulates distinct subsets of B cells, including naïve and memory B cells. Consistent with our findings, other studies have shown that MTX preferentially suppresses activated and memory B cell subsets, which are crucial for sustaining chronic inflammation and disease relapse in autoimmune conditions (46). This effect was not detected in patients receiving a biological DMARD (etanercept), which further highlights the role of MTX in modulating B cell subsets in autoimmune diseases (47) However, detailed studies analyzing B cell composition at different treatment time points are exceedingly rare. Collectively, our findings underscore the multifaceted role of MTX in modulating B cell subsets, thereby contributing to its therapeutic effects in autoimmune diseases.

Beyond its impact on B cells, we could detect that MTX targets RA-specific pathways specifically in key affected cell types such as T cells and monocytes. We identified a memory cluster predominantly composed of highly differentiated T cells (Fig. 1B). This cluster was more pronounced at baseline in RA patients compared to HC and we detected a clear regression within three weeks of MTX treatment. Within memory T cells, T follicular helper (Tfh) cells emerged as one of the most strongly modulated subsets by MTX. Tfh cells contribute to RA pathogenesis by promoting B cell proliferation, differentiation, and antibody maturation, as well as by expressing various immunomodulatory molecules (48–50). Consequently, reducing autoantibody production and restraining excessive Tfh cell responses are critical strategies for controlling RA pathogenesis. As highlighted in a recent review, Tfh cells play a central role in RA pathogenesis and represent promising therapeutic targets (51). In our study, transcriptional analysis of altered populations revealed pronounced modulations in IFN-α signaling, splicing, and cell cycle pathways within naïve and memory CD4^+^ T cells. The expression and function of associated apoptosis genes are altered in RA patients.

In the myeloid cluster, pDCs, cDCs, and monocytes were elevated in HC compared to RA baseline samples. Following MTX treatment, these populations increased in RA patients, reaching levels comparable to HC. Transcriptional analysis in our study revealed that monocytes were enriched in pathways related to autoimmunity, antigen processing, TNF-α, and IFN-γ responses. Consistent with this, studies examining the mode of action of MTX in monocytic cell lines have demonstrated its effects on the production of interleukin (IL)-1, IL-6, and TNF-α. Additionally, MTX modulated the expression of IL-1 and IL-6 as well as components of the Jun-N-terminal kinase (JNK) signaling pathway, including JNK1, JNK2, JUN, and FOS, and induced cell death in a dose-dependent manner *in vitro* (52). MTX appears to preferentially affect pro-inflammatory subsets, such as intermediate monocytes, which are known to produce higher levels of TNF-α and IL-6. This selective suppression modulates the inflammatory milieu without fully compromising innate immune function. Monocytes displayed enrichment in autoimmune and antigen processing pathways along with TNF-α and IFN-γ responses, whereas naive and memory CD4^+^ T cells were marked by pronounced modulations in IFN-α, splicing, and cell cycle pathways. Such specificity underscores prior reports that monocytes and T cells contribute distinctly to RA progression and demonstrates how MTX can exert wide-ranging yet cell-type-specific immunomodulatory actions (43, 53). Furthermore, gene regulatory network (GRN) analyses in monocytes and CD4^+^ memory T cells underscored the ability of MTX to restore a healthy-like transcriptional profile. Identified hub genes, including those previously implicated in RA pathogenesis (e.g., EGR1, JAK2, CXCL8, FCN1, SOCS1), exhibited restored expression patterns under MTX therapy, reflecting known immunoregulatory mechanisms (35–37, 41). These results are consistent with prior studies indicating that MTX mitigates pathogenic immune signaling by modulating key transcription factors (e.g., TNFAIP3, NFKBIZ) and immune-related pathways (54, 55). Novel targets, including ATP2B1-AS1, IGSF6, and SCO2 and members of the GIMAP family, warrant further exploration as potential contributors to RA pathogenesis and therapeutic targets. Altogether, the observed time-resolved shift toward a healthier gene regulatory architecture provides further evidence that MTX effectively rebalances aberrant immune responses in RA, reinforcing its status as a cornerstone therapy for this disease.

We acknowledge the limitations associated with the small sample size in this study, which imposes statistical constraints on our findings. However, the availability of high-quality patient samples over extended periods and multiple time points, as well as the accessibility of advanced methods remain limited to date. Clinical long-term follow up of the patients could provide valuable insights into temporal variations, facilitating the identification of permanently imprinted cell dynamics in contrast to early response dynamics.

In conclusion, this study demonstrates that methotrexate elicits early and cell-type-specific immunomodulatory effects in RA, particularly on T cells and monocytes. By employing an integrative multi-omics approach, we reveal that MTX rapidly reconfigures immune cell distributions and gene expression profiles toward a more “healthy-like” state. Despite the named limitations, these findings provide important insights into MTX’s mode of action and provide the potential to define early cellular biomarkers which can be tested in a prospective clinical trial setting.

## Data availability statement

Raw and processed data will be available upon acceptance. A cell browser website will be available to visualize our data and results.

## Acknowledgments

We thank all the patients who participated. We thank Martina Durechova, Daffodil Dioso and Michael Zauner for their support and Andreas Spittler, Brigitte Meyer, Carl-Walter Steiner, Birgit Niederreiter and Karolina von Dalwigk for their technical assistance. We thank Sylvia Taxer for her support and Birgit Reiter for helpful discussions on MTX metabolites. This study was supported by the MedUni Vienna Biobank KIP.

