## SupplementaryFigures for "Time-resolved immune dynamics in rheumatoid arthritis under methotrexate therapy"

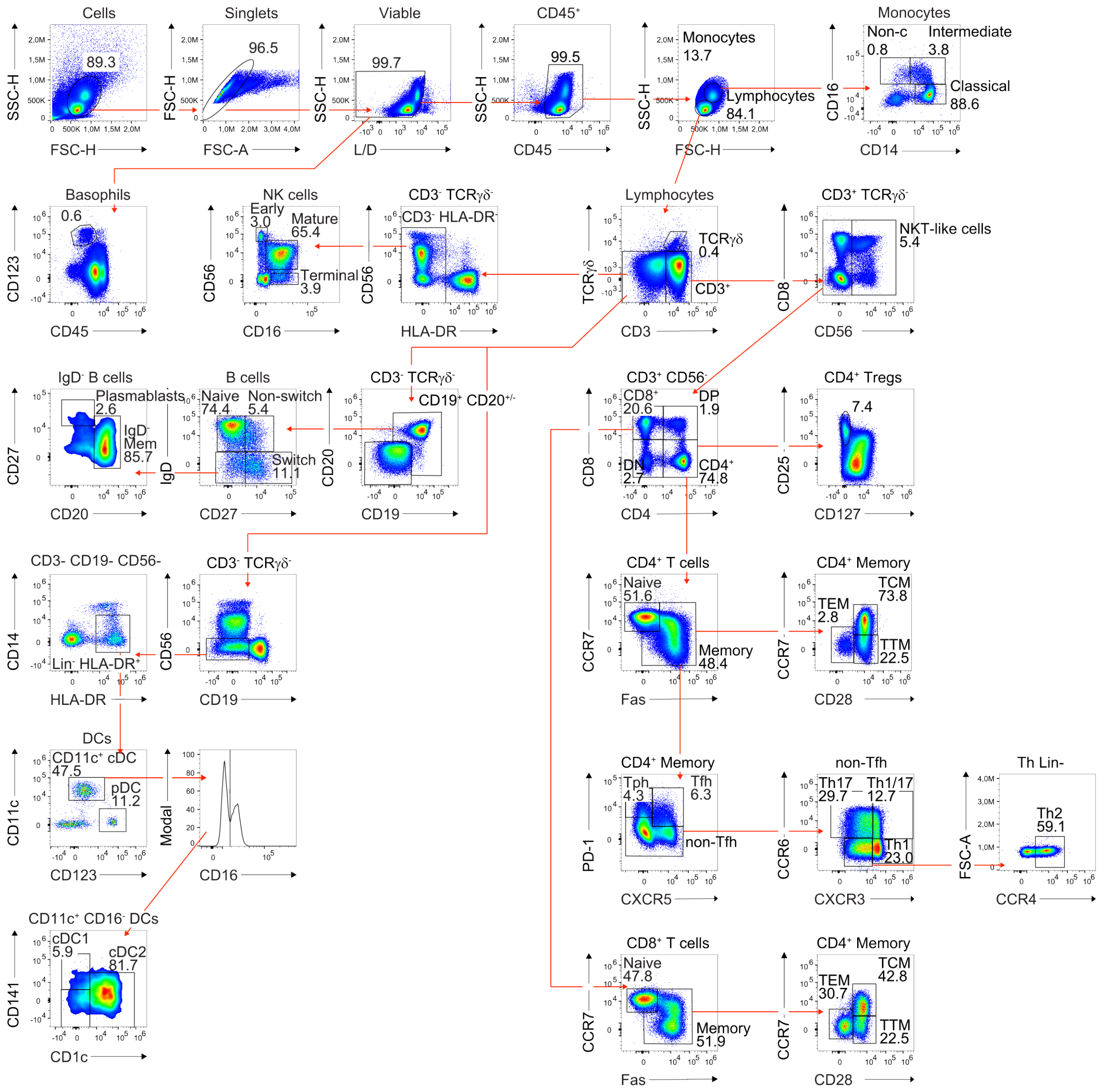


**Supplementary Figure 1. Manual gating strategy.** The gating strategy used to define major cellular subsets is illustrated, with arrows indicating relationships across plots and numerical annotations highlighting populations described in this analysis. After excluding doublets and dead cells, basophils were identified as $CD45^{+}CD123^{+}$ cells. Lymphocytes and monocytes were distinguished based on FSC-A/SSC-A properties. Monocytes were categorized by CD14 and CD16 expression into classical ($CD14^{+}CD16^{-}$), intermediate ($CD14^{+}CD16^{+/low}$), and non-classical ($CD14^{-}CD16^{+}$) subsets. From the lymphocyte gate, subsets including $CD3^{-}TCR\gamma\delta^{-}$, $CD3^{+}TCR\gamma\delta^{-}$, and $CD3^{+}TCR\gamma\delta^{+}$ cells were identified. Within the $CD3^{+}TCR\gamma\delta^{-}$ population, NKT-like ($CD3^{+}CD56^{+}$) subsets were delineated. $CD4^{+}$, $CD8^{+}$, $CD4^{+}CD8+$, and $CD4^{-}CD8^{-}$ T cells were gated from the $CD3^{+}CD56^{-}$ subset, with Tregs identified using $CD127^{low/-}CD25^{high}$ expression. The memory/effector states of $CD4^{+}$ and $CD8^{+}$ T cells were classified using CCR7, Fas, and CD28. T helper cell subsets were identified by the expression of

PD-1, CXCR5, CCR6, CCR4, and CXCR3. B cells were gated from the $CD3^{-}TCR\gamma\delta^{-}$ population as $CD19^{+}$ and/or $CD20^{+}$ cells and subtyped into $IgD^{+}CD27^{-}$, $IgD^{+}CD27^{+}$, and $IgD^{-}CD27^{+/-}$ populations; plasmablasts and $IgD^{-}$ memory B cells were distinguished by the expression of CD20 and CD27. NK cells were defined as $CD3^{-}TCR\gamma\delta^{-}HLA\text{-}DR^{-}$ cells and classified into early ($CD56^{+}CD16^{-}$), mature ($CD56^{+}CD16^{+}$), and terminal ($CD56^{-}CD16^{+}$) NK cells. Dendritic cells (DCs) were identified as $CD3^{-}CD19^{-}CD56^{-}CD14^{-}HLA\text{-}DR^{+}$ cells and subtyped into plasmacytoid ($CD123^{+}$) and conventional (CD11c+) DCs, with further classification of $CD11c^{+}$ DCs based on CD16, CD1c, and CD141 expression. All data were generated using frozen PBMCs from a single healthy donor.

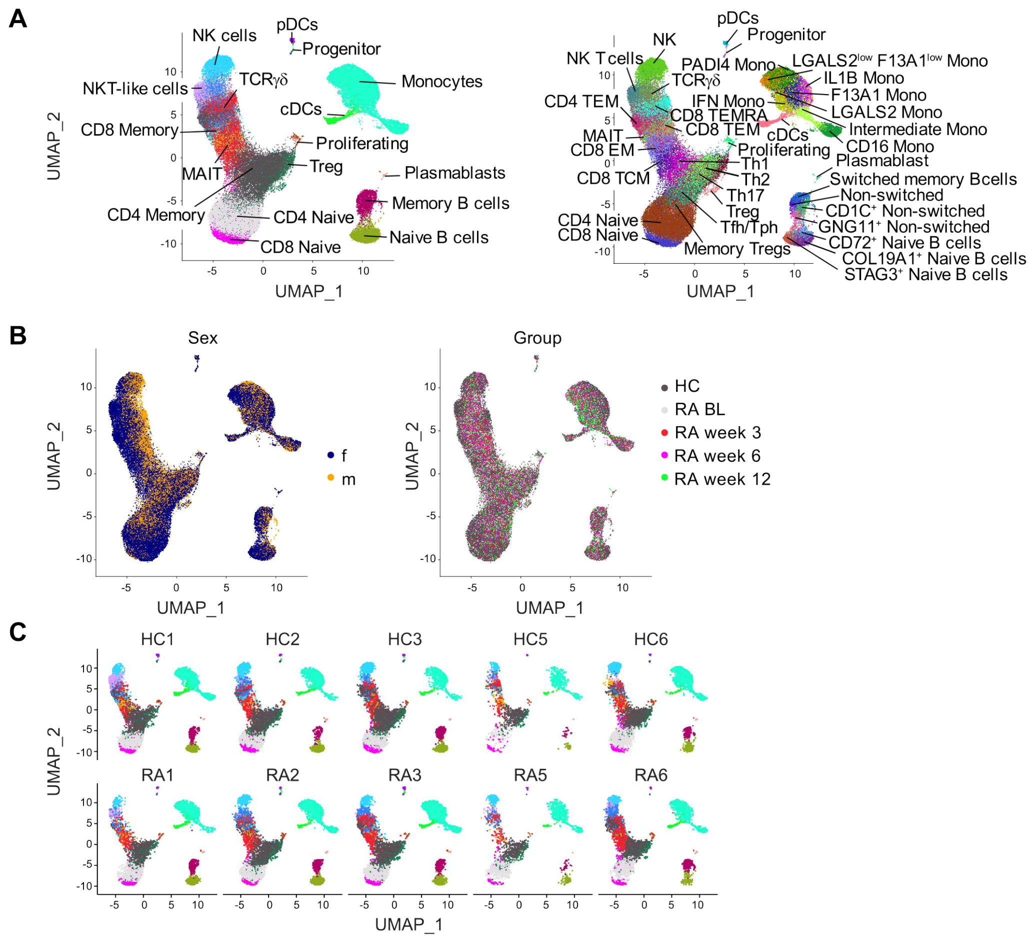


**Supplementary Figure 2. UMAPs showing the effect of covariates on scRNA-seq cluster distribution.** (A) Comparison of the course grained clusters (left panel) and fine grained clusters (right panel). (B) The effect of patient sex or group on the cluster distribution. (C) Cluster distribution per donor.

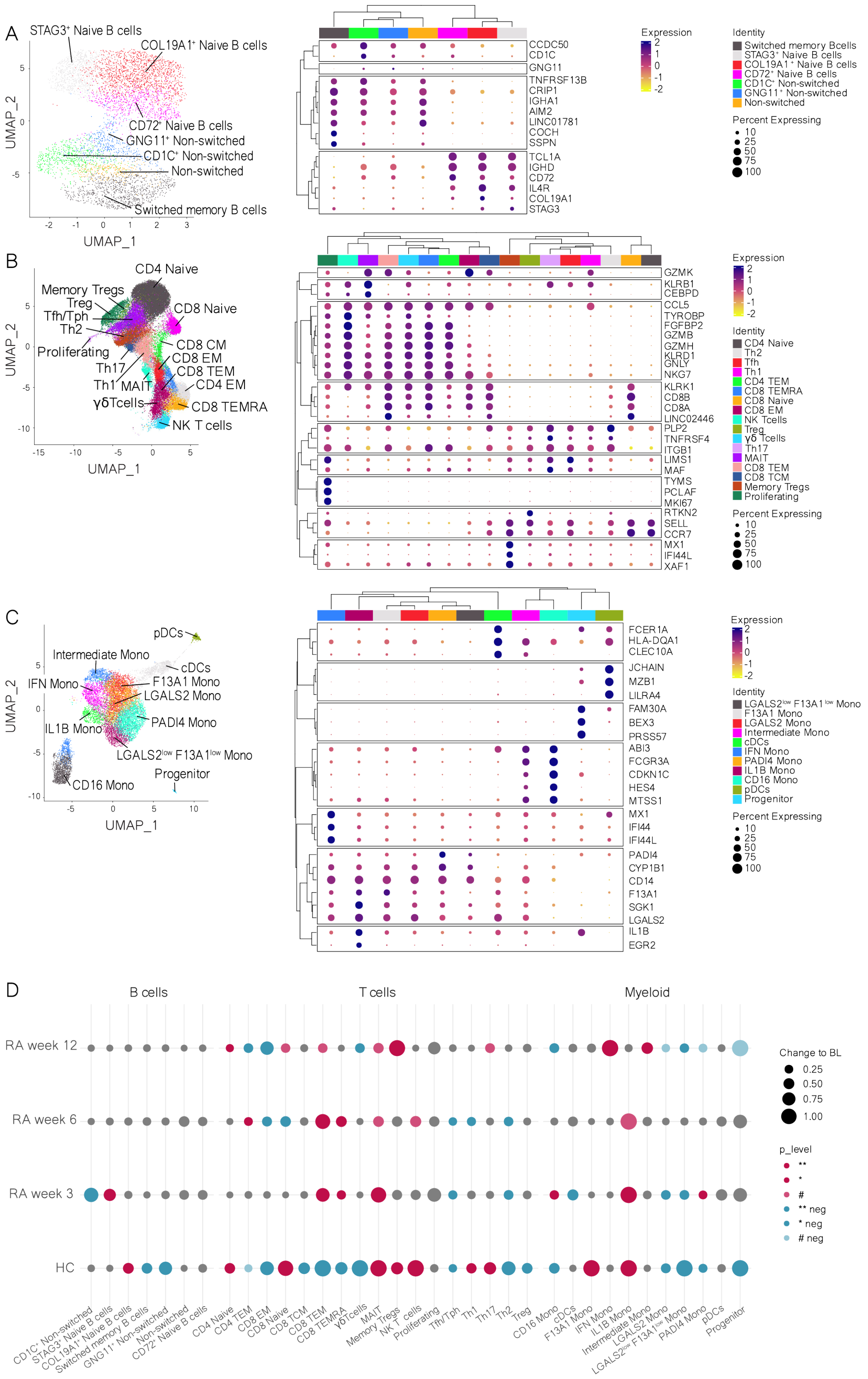
A
STAG3+ Naive B cells
COL19A1+ Naive B cells
CD72+ Naive B cells
GNG11+ Non-switched
CD1C+ Non-switched
Non-switched
Switched memory B cells
UMAP_2
UMAP_1
CCDC50
CD1C
GNG11
TNFRSF13B
CRIP1
IGHA1
AIM2
LINC01781
COCH
SSPN
TCL1A
IGHD
CD72
IL4R
COL19A1
STAG3
Expression
Identity
Switched memory Bcells
STAG3+ Naive B cells
COL19A1+ Naive B cells
CD72+ Naive B cells
CD1C+ Non-switched
GNG11+ Non-switched
Non-switched
Percent Expressing
B
CD4 Naive
Memory Tregs
Treg
Tfh/Tph
Th2
CD8 Naive
Proliferating
CD8 CM
CD8 EM
CD8 TEM
Th17
Th1
MAIT
CD4 EM
γδTcells
CD8 TEMRA
NK T cells
GZMK
KLRB1
CEBPD
CCL5
TYROBP
FGFBP2
GZMB
GZMH
KLRD1
GNLY
NKG7
KLRK1
CD8B
CD8A
LINC02446
PLP2
TNFRSF4
ITGB1
LIMS1
MAF
TYMS
PCLAF
MKI67
RTKN2
SELL
CCR7
MX1
IFI44L
XAF1
CD4 Naive
Th2
Tfh
Th1
CD4 TEM
CD8 TEMRA
CD8 Naive
CD8 EM
NK Tcells
Treg
γδ Tcells
Th17
MAIT
CD8 TEM
CD8 TCM
Memory Tregs
Proliferating
C
pDCs
Intermediate Mono
cDCs
IFN Mono
F13A1 Mono
LGALS2 Mono
PADI4 Mono
IL1B Mono
LGALS2low F13A1low Mono
Progenitor
CD16 Mono
FCER1A
HLA-DQA1
CLEC10A
JCHAIN
MZB1
LILRA4
FAM30A
BEX3
PRSS57
ABI3
FCGR3A
CDKN1C
HES4
MTSS1
MX1
IFI44
IFI44L
PADI4
CYP1B1
CD14
F13A1
SGK1
LGALS2
IL1B
EGR2
LGALS2low F13A1low Mono
F13A1 Mono
LGALS2 Mono
Intermediate Mono
cDCs
IFN Mono
PADI4 Mono
IL1B Mono
CD16 Mono
pDCs
Progenitor
D
B cells
T cells
Myeloid
RA week 12
RA week 6
RA week 3
HC
Change to BL
0.25
0.50
0.75
1.00
p_level
**
*
#
** neg
* neg
### neg
CD1C+ Non-switched
STAG3+ Naive B cells
COL19A1+ Naive B cells
Switched memory B cells
GNG11+ Non-switched
Non-switched
CD72+ Naive B cells
CD4 Naive
CD4 TEM
CD8 EM
CD8 Naive
CD8 TCM
CD8 TEM
CD8 TEMRA
γδTcells
MAIT
Memory Tregs
NK T cells
Proliferating
Tfh/Tph
Th1
Th17
Th2
Treg
CD16 Mono
cDCs
F13A1 Mono
IFN Mono
IL1B Mono
Intermediate Mono
LGALS2 Mono
LGALS2low F13A1low Mono
PADI4 Mono
pDCs
Progenitor

**Supplementary Figure 3. Fine grained-subset cluster distribution.** (A) UMAP plot showing fine-grained sub-clustering of B cells (left), and expression of the most prominent marker genes for each sub-cluster (right). (B) UMAP plot showing fine-grained sub-clustering of T cells (left), and expression of the most prominent marker genes for each sub-cluster (right). (C) UMAP plot showing fine-grained sub-clustering of myeloid cells (left), and expression of the most prominent marker genes for each sub-cluster (right). (D) Cell type abundance comparison across fine-grained cell subsets. Healthy controls and each MTX treatment timepoint (3, 6 and 12 weeks post MTX) were compared to baseline RA (BL). Cell types with an increase in abundance compared to BL are depicted in red, whereas decrease in blue. The size of the dots corresponds to the absolute observed log2 fold difference (log2FD) compared to BL, whereas the color intensity represents the adjusted p-value using a permutation test. Gray represents non-significant changes. The log2FD values are capped at 1. Significance thresholds were defined as follows: #P < 0.1, *P < 0.05, **P < 0.01, ***P < 0.001, ****P < 0.0001. Abbreviations: healthy controls (HC), rheumatoid arthritis (RA), baseline (BL), Natural Killer T cell (NKT) – like cells, Methotrexate (MTX), gamma-delta T cell (TCRgd), conventional Dendritic Cells (cDC), plasmacytoid Dendritic Cells (pDC), effector memory (EM), terminal effector memory (TEM), Mucosa-associated invariant T (MAIT), effector memory *cells* re-expressing CD45RA (TEMRA),  central memory (TCM).

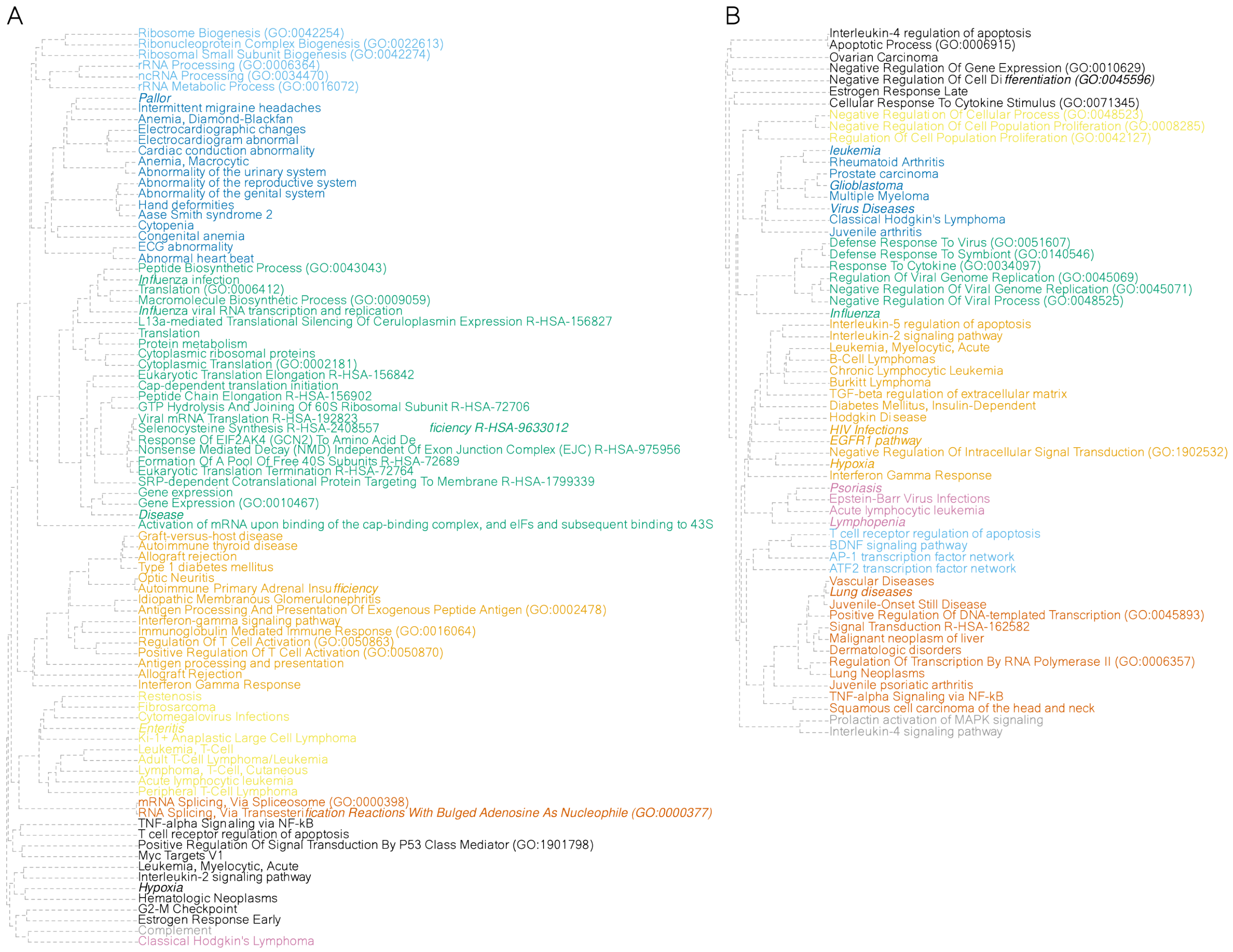


**Supplementary Figure 4. Dendrograms of associated terms corresponding to figures 3A and 3B.** (A) Dendrograms of associated terms enriched for differentially expressed genes between HC and RA BL. (B) Dendrograms of associated terms enriched for differentially expressed genes affected by MTX treatment.

A Regulatory networks - Monocytes

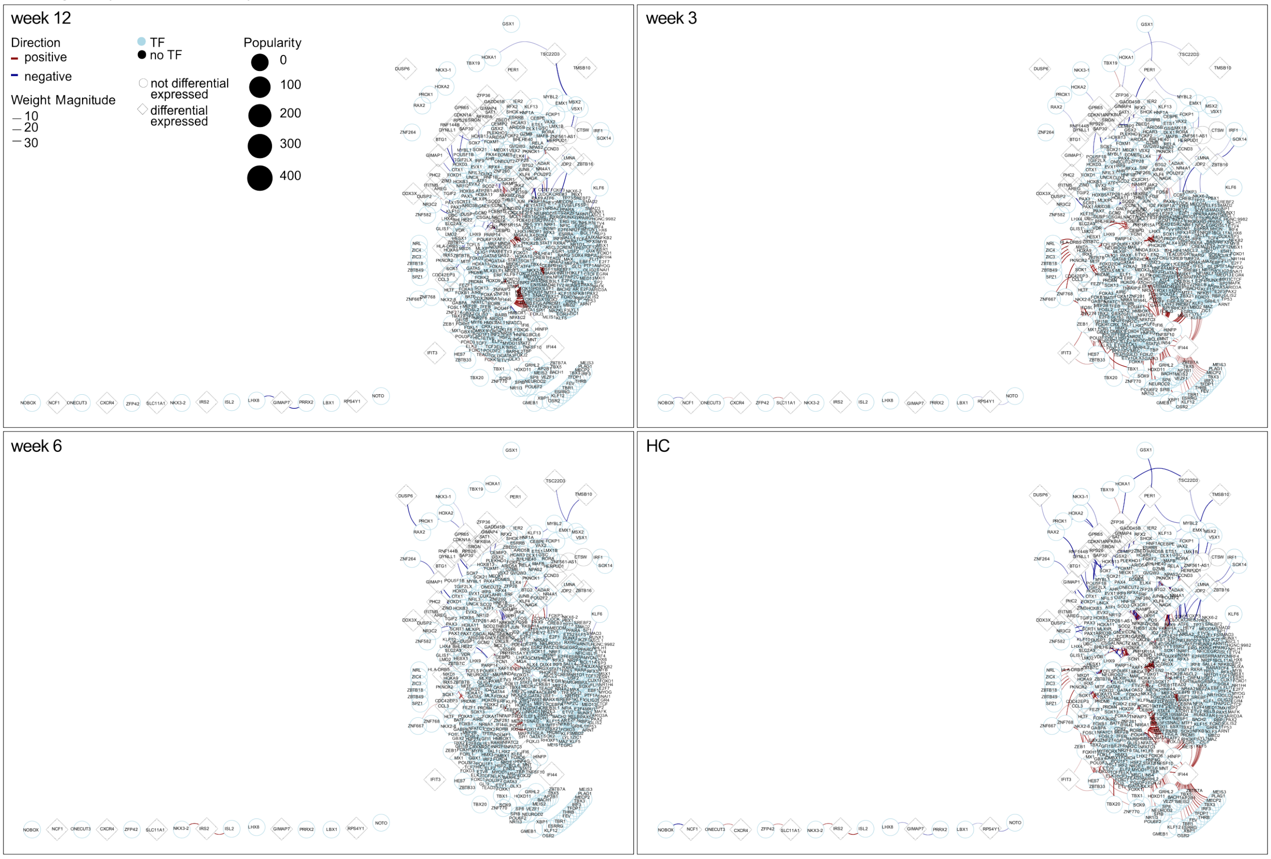


B Co-regulatory networks - Monocytes

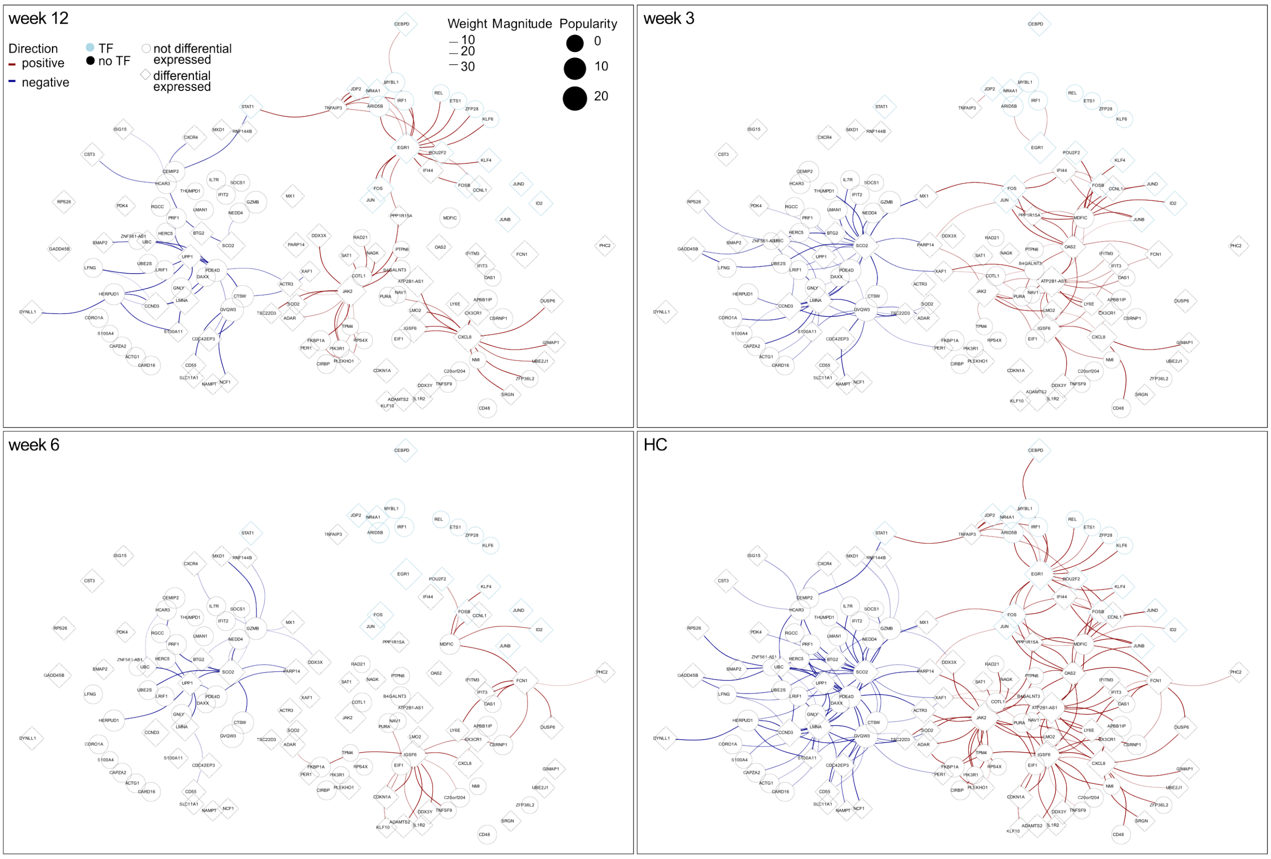

**Supplementary Figure 5. Extended gene regulatory networks for monocytes.** (A) Full regulatory network in monocytes; edges depict difference in co-regulation weights of two genes, one of which is a transcription factor, between either HC and RA BL or weeks 3, 6, and 12 of MTX treatment and RA BL, with red being a weaker relationship and blue stronger. The intensity of the color represents the magnitude of the difference.  Light blue nodes represent transcription factors and diamond-shaped nodes represent genes affected by MTX treatment in the corresponding cell type. (B) Full co-regulatory network in monocytes; edges depict the difference in co-expression weights between either HC and RA BL or weeks 3, 6, and 12 of MTX treatment and RA BL, with red being a weaker relationship and blue stronger. The intensity of the color represents the magnitude of the difference. Light blue nodes represent transcription factors and diamond-shaped nodes represent genes affected by MTX treatment in the corresponding cell type.

A Regulatory networks - CD4 memory cells

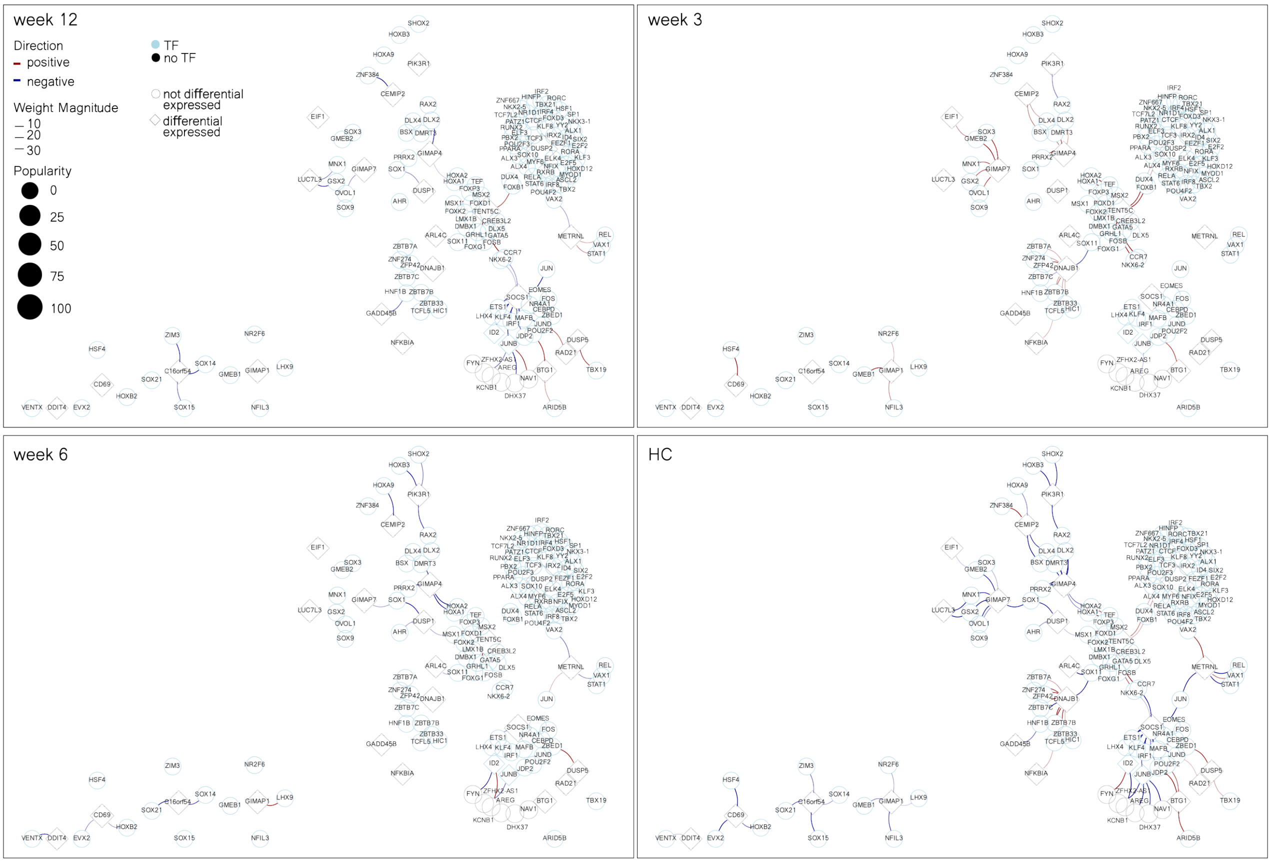


B Co-regulatory networks - CD4 memory cells

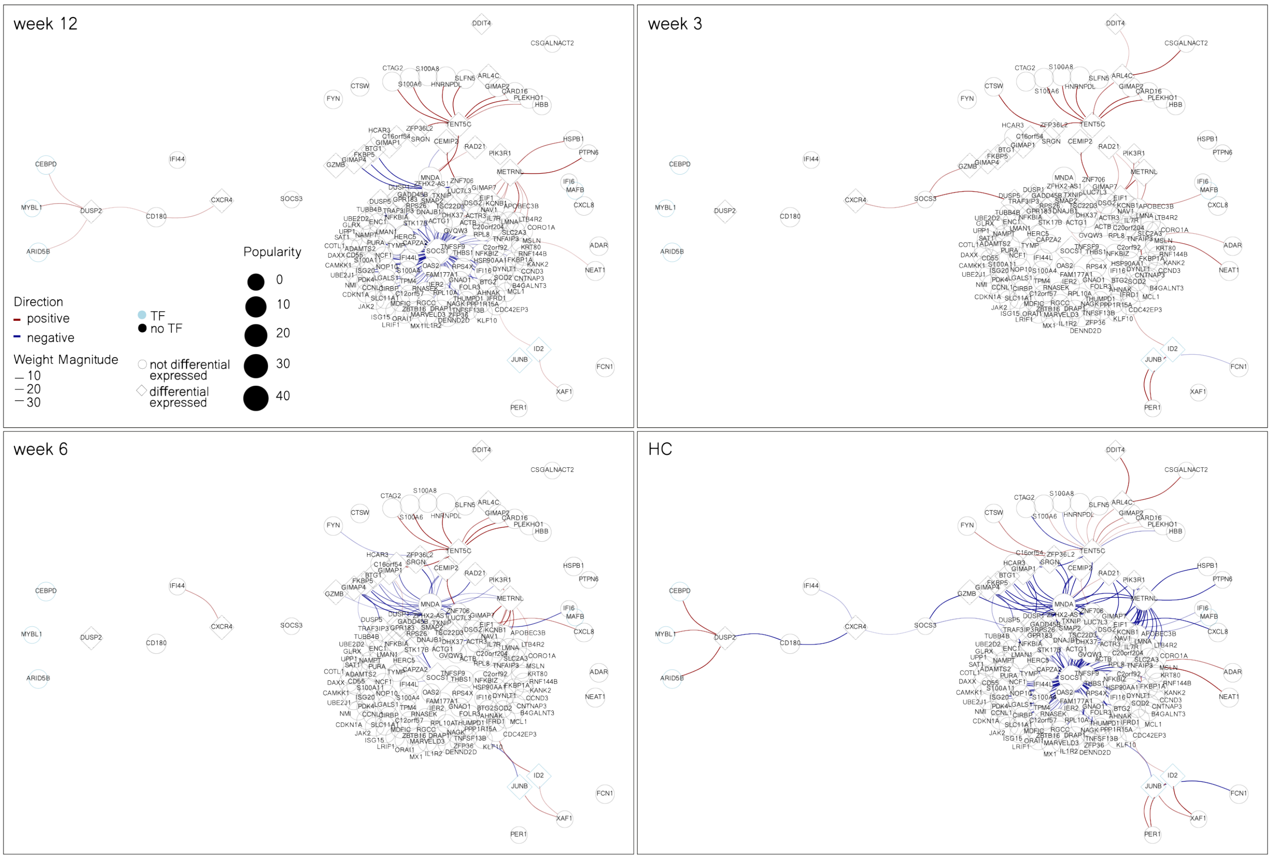

**Supplementary Figure 6. Extended gene regulatory networks for CD4 memory T cells.** (A) Full regulatory network in CD4 memory T cells; edges depict difference in co-regulation weights of two genes, one of which is a transcription factor, between either HC and RA BL or weeks 3, 6, and 12 of MTX treatment and RA BL, with red being a weaker relationship and blue stronger. The intensity of the color represents the magnitude of the difference. Light blue nodes represent transcription factors and diamond-shaped nodes represent genes affected by MTX treatment in the corresponding cell type. (B) Full co-regulatory network in CD4 memory T cells; edges depict the difference in co-expression weights between either HC and RA BL or weeks 3, 6, and 12 of MTX treatment and RA BL, with red being a weaker relationship and blue stronger. The intensity of the color represents the magnitude of the difference. Light blue nodes represent transcription factors and diamond-shaped nodes represent genes affected by MTX treatment in the corresponding cell type.

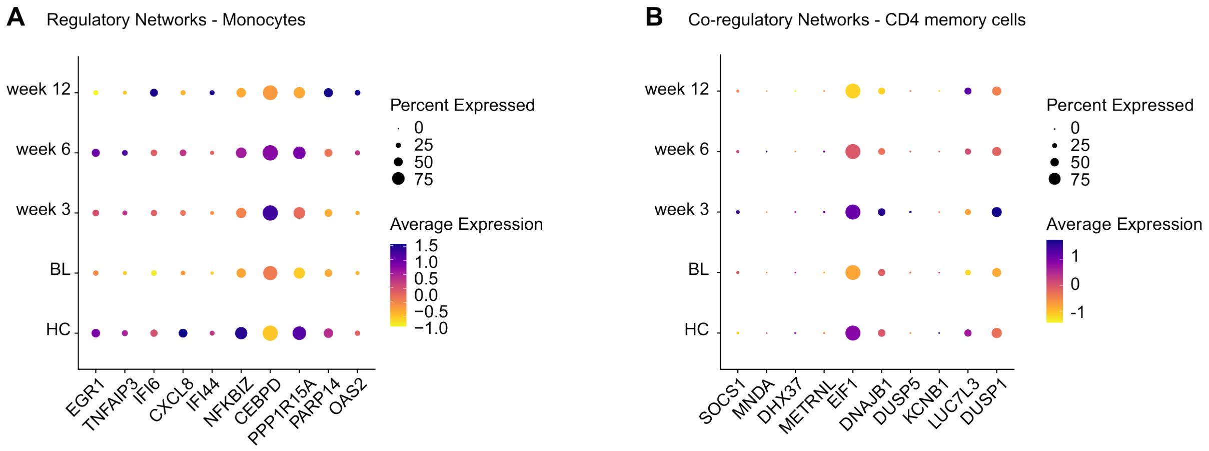


**Supplementary Figure 7. Groupwise expression of hub genes.** (A) Hub genes corresponding to supplementary figure 5A. Color depicts the scaled average expression of the gene in monocytes across groups. Dot size represents the percentage of cells with non-zero expression of the gene. (B) Hub genes corresponding to supplementary figure 6B. Color depicts the scaled average expression of the gene in CD4 memory T cells across groups. Dot size represents the percentage of cells with non-zero expression of the gene.
